# AP-1 Imprints a Reversible Transcriptional Program of Senescent Cells

**DOI:** 10.1101/633594

**Authors:** Ricardo Iván Martínez-Zamudio, Pierre-François Roux, José Américo N L F de Freitas, Lucas Robinson, Gregory Doré, Bin Sun, Jesús Gil, Utz Herbig, Oliver Bischof

**Affiliations:** Institut Pasteur, Laboratory of Nuclear Organization and Oncogenesis, Department of Cell Biology and Infection, 75015 Paris, France; INSERM, U993, 75015 Paris, France; Equipe Labellisée Fondation ARC pour la recherche sur le cancer, Paris, France; Université de Paris, Sorbonne Paris Cité, Paris, France; MRC London Institute of Medical Sciences (LMS), Du Cane Road, London, W12 0NN, UK; Institute of Clinical Sciences (ICS), Faculty of Medicine, Imperial College London, Du Cane Road, London W12 0NN, UK; Department of Microbiology, Biochemistry & Molecular Genetics, Rutgers Biomedical & Health Sciences, Rutgers University, 205 South Orange Avenue, Newark, NJ 07103, USA

## Abstract

Senescent cells play important physiological- and pathophysiological roles in tumor suppression, tissue regeneration, and aging. While select genetic and epigenetic elements crucial for senescence induction were identified, the dynamics, underlying epigenetic mechanisms, and regulatory networks defining senescence competence, induction and maintenance remain poorly understood, precluding a deliberate therapeutic manipulation of these dynamic processes. Here, we show, using dynamic analyses of transcriptome and epigenome profiles, that the epigenetic state of enhancers predetermines their sequential activation during senescence. We demonstrate that activator protein 1 (AP-1) ‘imprints’ the senescence enhancer landscape effectively regulating transcriptional activities pertinent to the timely execution of the senescence program. We define and validate a hierarchical transcription factor (TF) network model and demonstrate its effectiveness for the design of senescence reprogramming experiments. Together, our findings define the dynamic nature and organizational principles of gene-regulatory elements driving the senescence program and reveal promising inroads for therapeutic manipulation of senescent cells.

## INTRODUCTION

Cellular senescence plays beneficial roles during embryonic development, wound healing, and tumor suppression. Paradoxically, it is also considered a significant contributor to aging and age-related diseases including cancer and degenerative pathologies^1^. As such, research on therapeutic strategies exploiting senescence targeting (*e.g.*, senolytics, senomorphics or pro-senescence cancer therapies) to improve healthspan has gained enormous momentum in recent years^2^.

Cellular senescence is a cell fate that stably arrests proliferation of damaged and dysfunctional cells as a complex stress response. The most prominent inducers of senescence are hyper-activated oncogenes (oncogene-induced senescence, OIS)^3^. The senescence arrest is accompanied by widespread changes in gene expression, including a senescence-associated secretory phenotype (SASP) – the expression and secretion of inflammatory cytokines, growth factors, proteases, and other molecules, which exert pleiotropic effects on senescent cells themselves as well as the surrounding tissue^4^. Importantly, although activation of the senescence program can pre-empt the initiation of cancer, the long-term effects of the SASP make the local tissue environment more vulnerable to the spread of cancer and other age-related diseases. Therefore, therapeutic interventions aimed at limiting SASP production are of relevance for cancer and many age-related diseases^4–6^.

The knowledge on epigenetic mechanisms underlying senescence has only recently increased revealing significant contributions of select transcription factors (TFs), chromatin modifiers and structural components, as well as non-coding RNAs to the senescent phenotype^7–12^. A major limitation of such studies, however, was their restriction to start-end-point comparisons, ignoring the dynamic nature of the senescence fate transition. Consequently, critical gene-regulatory aspects of the execution and maintenance of the senescence state remain poorly understood. Therefore, an integrative, temporally resolved, multidimensional profiling approach is required to establish essential regulatory principles that govern this key biological decision-making process. Such knowledge would be instrumental both for identifying stage-specific senescence regulators and urgently needed specific biomarkers as well as control points in TF and gene regulatory networks, which would pave the way for a deliberate therapeutic manipulation of the senescence cell fate.

Enhancers are key genomic regions that drive cell fate transitions. The enhancer landscape is established during development by the concerted action of TF networks and chromatin modifiers^13^. The details on how this information converges in *cis* remain unclear, and we still lack valid organizational principles that explain the function of mammalian TF networks. In mammalian cells, enhancer elements are broadly divided into two major categories-active and poised. While active enhancers are characterized by the simultaneous presence of H3K4me1 together with H3K27ac and are associated with actively transcribed genes, only H3K4me1 marks poised enhancers, and their target genes are generally not expressed^14^. A subset of enhancers may also be activated *de novo* from genomic areas devoid of any TF binding and histone modifications. These latent or nascent enhancers serve an adaptive role in mediating stronger and faster gene expression upon cycles of repeated stimulation^15,16^. Recent evidence showed a role for enhancer remodeling in driving senescence-associated gene expression^11,12,17^. However, it is currently unknown which enhancer elements, epigenetic marks or TFs render cells competent to respond to senescence-inducing signals with precise timing and output. A thorough understanding of how senescence competence is established, realized and what defines it would allow for the prediction of a positive senescence engagement for example in pro-senescence cancer therapies^18^.

Pioneer TFs are critical in establishing new cell fate competence by granting long-term chromatin access to non-pioneer factors and are also crucial determinants of cell identity through their opening and licensing of the enhancer landscape^19,20^. We can now reliably deduce pioneer and non-pioneer TF activity from chromatin accessibility data allowing for the hierarchization of TF function whereby pioneer TFs sit atop a TF binding hierarchy, recruiting non-pioneers such as settler and migrant TFs to gene-regulatory regions for optimal transcriptional output^21^. The pioneer TFs bestowing senescence potential have not been identified to date. However, their identification might be a pre-requisite for reprogramming or manipulation of senescent cells for future therapeutic benefit as was shown successfully for the reprogramming to adopt full stem cell identity^22^.

In this study, we examined the possibility that the epigenetic state of enhancers could determine senescence cell fate. We explored this question by generating time-resolved transcriptomes and comprehensive epigenome profiles during oncogenic RAS-induced senescence. Through integrative analysis and further functional validation, we revealed novel and unexpected links between enhancer chromatin, TF recruitment, and senescence potential and defined the organizational principles of the TF network that drive the senescence program. Together, this allowed us to precisely manipulate the senescence phenotype with important therapeutic implications. Specifically, we show that the senescence program is predominantly encoded at the enhancer level and that the enhancer landscape is dynamically reshaped at each step of the senescence transition. Remarkably, we find that this process is pre-determined before senescence induction and AP-1 acts as a pioneer TF that ‘premarks’ prospective senescence enhancers to direct and localize the recruitment of other transcription factors into a hierarchical TF network that drives the senescence transcriptional program after induction. We also uncover a class of enhancers that retain an epigenomic memory after their inactivation during the senescence transition. These “remnant” enhancers lack traditional enhancer histone-modification marks but are instead “remembered” by AP-1 TF bookmarking for eventual future re-activation. Finally, functional perturbation of prospective senescence enhancers and AP-1 validated and underscored the importance of these entities for the timely execution of the senescence gene expression program and allowed for the precise reprogramming and reversal of the senescence cell fate.

## RESULTS

We employed time-series experiments on WI38 fibroblasts undergoing oncogene-induced senescence (OIS) using a tamoxifen-inducible ER: RASV12 expression system^23^. We determined global gene expression profiles by microarrays and mapped the full set of accessible chromatin sites by ATAC-seq^24^ at 6-time points (0, 24, 48, 72, 96 and 144 h). Cells intended for ChIP-seq were crosslinked at 3-time-points (0, 72 and 144h) and used for profiling histone modifications including H3K4me1 (putative enhancers), H3K4me3 (promoters), H3K27ac (active enhancers and promoters) and H3K27me3 (polycomb repressed chromatin). From accessible chromatin regions determined by ATAC-seq we deduced TF binding dynamics and hierarchies (Figure 1A). For comparison, we included cells undergoing quiescence (Q) by serum withdrawal for up to 96h. Unlike senescence arrested cells, quiescence arrested cells can be triggered to re-enter the cell cycle upon serum addition. Q and OIS cells were validated using classical markers (Supplementary Figures 1A-B).

**FIGURE 1:**
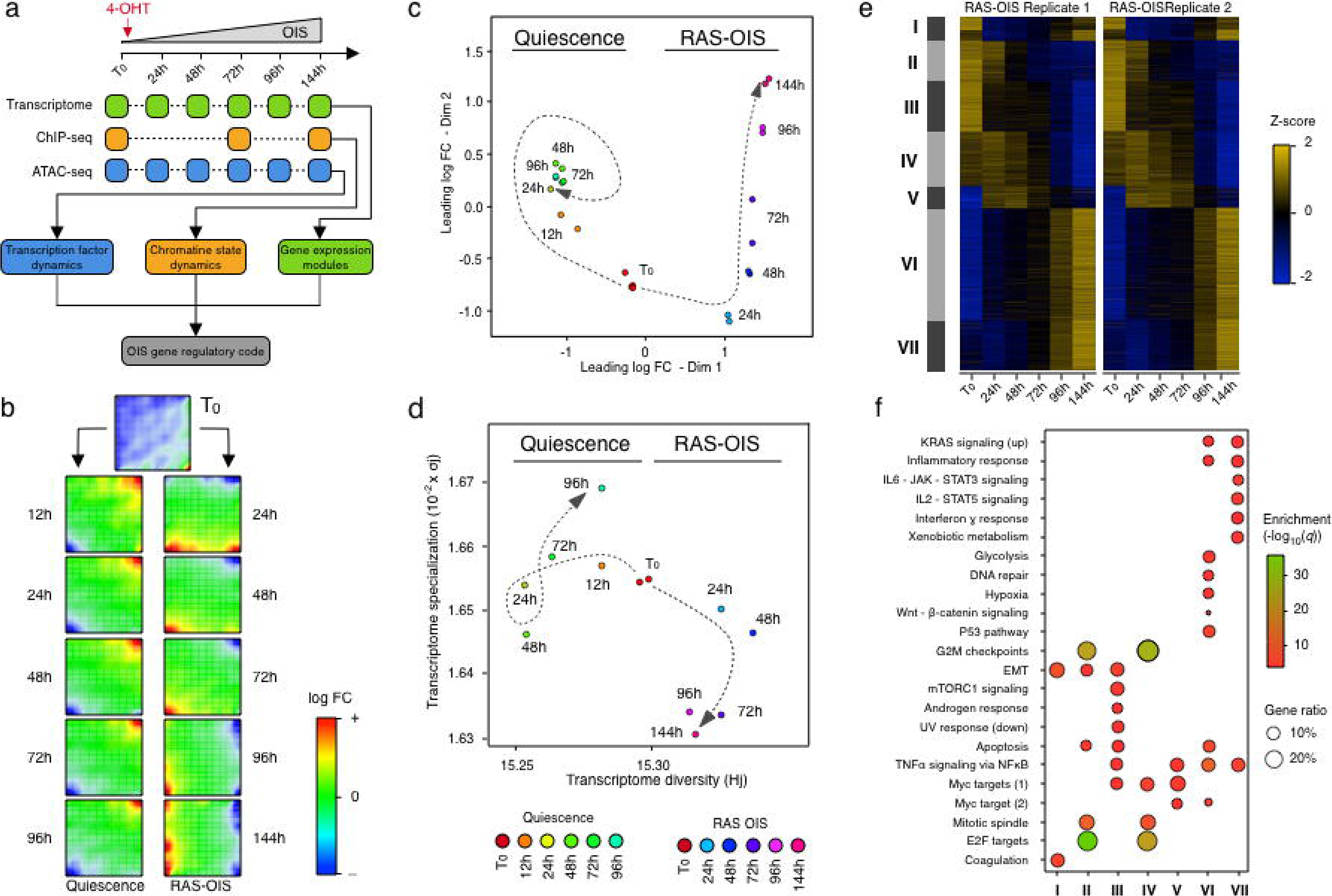
Multi-state establishment of the senescence transcriptional program. **(a)** Schematic overview for defining the gene-regulatory code of RAS-OIS using time-resolved, high-throughput transcriptome (microarray) and epigenome (ChIP-seq, and ATAC-seq) data sets. **(b)** Self-organizing maps (SOMs) of gene expression profiles for quiescence and RAS-OIS time-series experiments as logarithmic fold-change. Red spots mark overexpression, blue spots underexpression. **(c)** Multidimensional scaling (MDS) analysis scatter plot visualizing the level of similarity/dissimilarity between normalized quiescence and RAS-OIS time-series transcriptomes. Distances between samples represent leading logarithmic fold-changes defined as the root-mean-squared average of the logarithmic fold-changes for the genes best distinguishing each pair of samples. **(d)** Scatter plot depicting the evolution of transcriptome diversity (H_j_) *vs.* transcriptome specialization (σ_j_) in cells undergoing quiescence or RAS-OIS. For each time-point and treatment, the average H_j_ and σ_j_ values across biological replicates are given. T_0_ is start of time-course. **(e)** Heatmap showing seven modules (I-VII) of temporally co-expressed genes specific for RAS-OIS using an unsupervised WGCNA clustering approach. Data are expressed as raw Z-scores. **(f)** Functional over-representation map depicting Molecular Signatures Database (MSigDB) hallmark gene sets associated to each transcriptomic cluster. Dots are color-coded according to the FDR corrected *p*-value based on the hypergeometric distribution. Size is proportional to the percentage of genes in the gene set belonging to the cluster.

### Multi-state establishment of the senescence transcriptional program

To identify and visualize dynamic gene expression patterns across the entire Q and RAS-OIS time-courses, we employed an unsupervised, self-organizing map (SOM) machine learning technique^25^ (Figure 1B) and multidimensional scaling (MDS) (Figure 1C) to our transcriptome data sets. Remarkably, serum-deprived fibroblasts rapidly established a Q-specific gene expression program within 24 h after serum deprivation, which changed only marginally within the remainder of the time-course (Figure 1B, left column and Figure 1C), and mainly involves only up-regulated (Figure 1B, top right corner, red) and down-regulated (Figure 1B, bottom left corner, blue) genes. By contrast, fibroblasts undergoing RAS-OIS displayed dynamic gene expression trajectories that evolved steadily, both for up- (red) and down-regulated metagenes (blue) (Figures 1B, right column and Figure 1C). Thus, RAS-OIS, unlike Q, is highly dynamic and does not gyrate towards a stable transcriptome end state. To substantiate this further we calculated the diversity and specialization of transcriptomes and gene specificity^26^ (Figure 1D) and performed a kernel density estimation analysis (Supplementary Figure 1C). These analyses demonstrated that RAS-OIS cells exhibit a temporally evolving increase in transcriptional diversity, whereas Q cells exhibit a temporally evolving, specific gene expression program. We conclude that the RAS-OIS cell fate is an open-ended succession of cell states rather than a fixed cell fate with a defined end-point, which is the current tenet. The apparent open-endedness and transcriptional diversity may provide a fertile soil for the eventual escape of pre-cancerous senescent cells as previously shown^27,28^.

To further delineate the evolution of the RAS-OIS gene expression program, we next performed dynamic differential gene expression analysis on the Q and OIS datasets^29^. A total of 4,986 genes (corresponding to 2,931 up-regulated and 2,055 down-regulated genes) were differentially regulated in at least one-time point (with a minimal leading log2 fold-change of 1.2; q=5*10^−4^) and partitioned into seven (I-VII) gene expression modules with distinct functional overrepresentation profiles in line with the senescence phenotype (Figures 1E-F and Figure S1D). The highly reproducible dynamics of gene expression during RAS-OIS transition suggest a high degree of preprogramming of this succession of cell states.

Cell-fate decisions are typically associated with stable changes in gene expression that shift the regulatory system from one steady state to the next^30^. In line with this, we found that proliferation-promoting genes of modules II and IV (E2F targets and G2M checkpoint) became increasingly repressed (*i.e.* senescence arrest), while pro-senescent SASP genes of modules VI and VII (*e.g.*, inflammatory and interferon response genes) became persistently induced (Figure 1F and Supplementary Figure 1E). Apoptosis-related genes of module III were repressed very early on in the time-course (within the first 24-48 hours during RAS-OIS induction), which is surprising given that apoptosis-resistance is considered a very late event in senescence (Figure 1F and Supplementary Figure 1E). This indicates that the commitment to senescence is a very early event made at the expense of apoptosis. Finally, we identified a set of genes in modules I and V that would have gone unnoticed in a traditional start-end-point analysis because they follow an “impulse”-like pattern. In these modules, transcript levels spiked-up (module V) or down (module I) following RAS-OIS induction, then sustained a new level, before transitioning to a new steady state, similar to the original levels (Figures 1E and Supplementary Figure 1E). These expression patterns support the notion that genes of module V play an active and vital role early in the transition to RAS-OIS, while genes in module I hold key regulators to maintain the proliferative fibroblast state.

Altogether, our investigation of transcriptome dynamics defined a modular organization and transcriptional diversity of the RAS-OIS gene expression program, providing a framework to unravel the gene-regulatory code underlying the senescence process.

### A dynamic enhancer program shapes the senescence transcriptome

Senescence cell fate involves a global remodeling of chromatin and specifically, the enhancer landscape^11,12^. An unanswered question, however, is how TFs and epigenetic modifications cooperatively shape a transcriptionally permissive enhancer landscape prior to gene activation to endow the cell with senescence potential.

To provide mechanistic insight into this question, we first comprehensively mapped genomic regulatory elements (*i.e.* putative enhancers, promoters and polycomb-repressed chromatin) during the transition of proliferating cells to RAS-OIS, profiling genome-wide histone modifications by ChIP-seq and transposon-accessible chromatin by ATAC-seq. To capture and quantify chromatin state dynamics we used ChromstaR (see Materials and Methods), which identified a total of sixteen chromatin states (Supplementary Figure 2A). The majority of the genome (≈80%) was, irrespective of the time-point, either devoid of any of the histone modifications analyzed (≈62%) or polycomb-repressed (≈18%). The fraction of the genome represented by active and accessible chromatin states (*i.e.*, enhancers and promoters) was comparably lower (≈20% combined). Chromatin state transitions occurred most prominently at enhancers, while promoters were only mildly affected (Figures 2A-B and Supplementary Figure 2A, arrows) congruent with previous results^11^. Unexpectedly, we found, however, that most of the enhancer activation, *i.e.* acquisition of H3K4me1 and H3K27ac, occurred *de novo* from unmarked chromatin at the T0-72 h and 72 h-144 h intervals, followed by the more stereotypical enhancer activation from a poised state (H3K4me1^+^ plus H3K27ac acquisition) and enhancer poising from the unmarked and polycomb-repressed state at the T0-72 h interval (acquisition of H3K4me1) (Figures 2A and -B). Thus, the regulatory landscape of senescence is largely predetermined by sequential enhancer activation from *de novo* and poised enhancers implying the existence of a prior imprint of past cell fate decisions.

The (in)activation chronology of enhancers was highly concordant with the temporal expression pattern of the nearest genes, indicating that most of these elements indeed function as *bona fide* enhancers (Supplementary Figure 2B). In line with this, correspondence analysis (CA) (Supplementary Figure 2C) revealed a strong correlation between gene expression modules (Figure 1E) and chromatin state transitions (Figure 2A). For example, globally up-regulated transcriptomic modules V, VI, VII projected proximally to chromatin state transitions involving enhancer activation congruent with the installation of the SASP. By contrast, dynamic enhancer inactivation associated with repressed transcriptomic modules (II, III, IV) congruent with installation of the senescence arrest. Finally, the oscillatory expression of genes in the module I associated with an equally oscillatory activation of its closest enhancers. Therefore, dynamic remodeling of the enhancer landscape reflects and defines the modular and dynamic nature of the RAS-OIS gene expression program.

**FIGURE 2:**
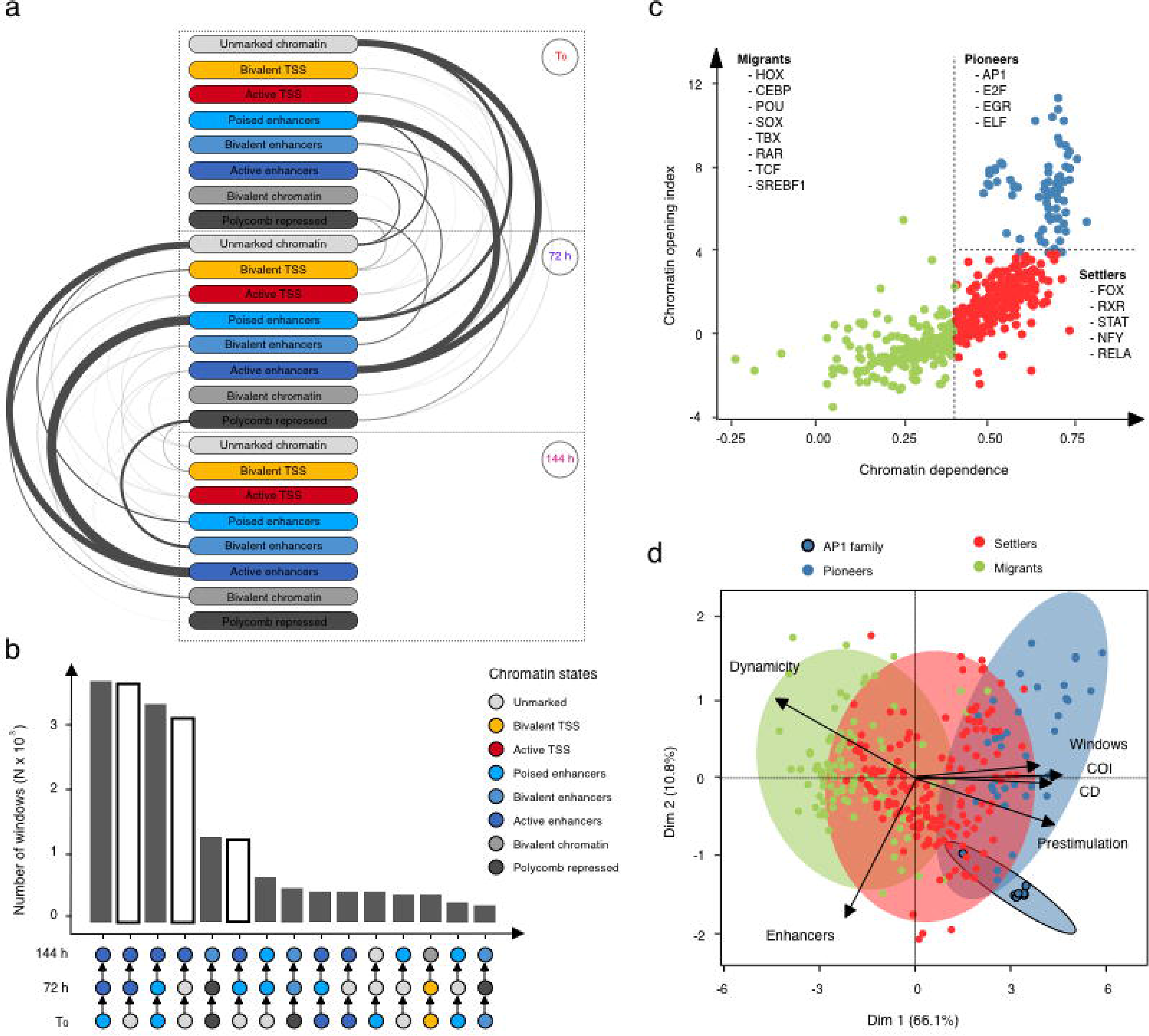
A dynamic enhancer program shapes the senescence transcriptome. **(a)** Arc plot visualizing dynamic chromatin state transitions for the indicated intervals. Edge width is proportional to the number of transitions. **(b)** Histogram showing the total number of windows of the top 15 chromatin states transitions. Chromatin state transitions corresponding to *de novo* enhancer activation are highlighted as white bars. **(c)** Chromatin dependence (CD) versus chromatin opening index (COI) are plotted for high-confidence TF sequence motifs used in our study (see Materials and Methods for details). Pioneer, settler and migrant TFs as defined by their COI and CD property are color-coded and select members of each TF class are listed. Same color code is used in all figures. **(d)** Biplot for principal component analysis performed with select TF binding parameters: dynamicity, total number of bound windows (N), percentage of binding at enhancers, pioneer index (referred to as the number of bound windows pre-stimulation), chromatin opening index (COI) and chromatin dependence (CD). The plot depicts the projections of the TFs and the loading of the different covariates for the first two principle components which explain 76.9% of the total inertia. The ellipses delineate the 95% confidence intervals for AP1 pioneers (blue with black outline), non-AP1 pioneers (blue), settlers (red), and migrants (green).

We next addressed the question of which TFs are key drivers for the dynamic enhancer remodeling driving the senescence transcriptome. To this end, we first intersected ATAC-seq peaks with the identified enhancer coordinates (Figure 2A-B) and performed a motif over-representation test. This analysis identified AP-1 super-family members (cJUN, FOS, FOSL1, FOSL2, BATF) as well as AP-1-associated TFs ATF3 and ETS1 as the most enriched motifs at any given time-point, thus, hinting at a putative chromatin priming and pioneer function for these TFs (Supplementary Figure 2D). Because AP-1 TFs are essential and inducible downstream effectors for the RAS signaling pathway in cellular transformation^31^ the possibility remains that the observed enrichment of AP-1 TFs at enhancers is strictly dependent on oncogenic RAS signaling *per se* and not a reflection of a specific pioneering role in the enhancer landscape independent of RAS signaling. We therefore performed the same analysis in cells undergoing replicative senescence (which is driven by loss of telomere integrity) and also in growth factor-deprived (and thus RAS signaling-muted) quiescent cells (Supplementary Figures 2E-F). In both cases, the AP-1 motif ascended as the predominant motif enriched at enhancers, thus, corroborating the notion that AP-1 TFs act as the universal pioneers imprinting the global as well as senescence-associated enhancer landscape.

To elaborate this further, we analyzed our time-resolved ATAC-seq data sets by adapting the “Protein Interaction Quantitation (PIQ)” algorithm, which was developed initially for DNAse-seq-based digital TF footprinting^21^. Importantly, PIQ allows for the functional hierarchization of TFs into pioneers, settlers, and migrants - whereby pioneer TFs bind nucleosome-compacted chromatin to initiate chromatin remodeling and to enable subsequent binding of non-pioneers (*i.e.*, settler and migrant TFs). PIQ segregated TFs into pioneers (*e.g.*, AP-1 TF family members), settlers (*e.g.*, NFY and RELA subunit of NF-κB) and migrants (*e.g.*, TF RAR family members and SREBF1) (Figure 2C). We confirmed this TF hierarchization by inspecting a selection of individual TF footprints for their adjacent nucleosomal positioning (Supplementary Figure 2G-I). AP-1 family member FOSL1, for example, bound to its cognate binding site despite the presence of strongly positioned flanking nucleosomes, as would be expected from a pioneer TF (Supplementary Figure 2G), while RELA binding required partial nucleosome displacement/chromatin opening, as would be expected for a settler TF (Supplementary Figure 2H), and SREBF1 bound to its cognate site in a near-nucleosome free context, as would be expected for a migrant TF (Supplementary Figure 2I). Importantly, there was a high correspondence between PIQ predictions and TF ChIP-seq profiling as exemplified for AP-1-members FOSL2 and cJUN, which we used as surrogate marks for bound AP-1 (which is typically a complex of JUN-JUN or JUN-FOS family member dimers), and RELA (Supplementary Figure 2J).

To decode additional TF properties critical for shaping the dynamic RAS-OIS enhancer landscape, we applied a principal component analysis (PCA) considering several metrics describing TF binding characteristics (Figure 2D). This analysis revealed two key features: First, pioneer TFs bind statically, extensively, and most importantly before RAS-OIS induction (*i.e.*, pre-stimulation) along the genome, while settler and migrant TFs bind more dynamically (“Dynamicity” in Figure 2D), far less frequently (“Windows” in Figure 2D), and on average less often before OIS induction (*i.e.* pre-stimulation) along the genome. Second, and in line with the proposed pioneering activity of AP-1 TFs, the latter clearly stand out amongst other pioneer TFs (highlighted by black circle in Figure 2D) because they bind exclusively and extensively to enhancers prior to RAS-OIS induction whereas most of the remaining pioneer TFs tend to accumulate away from them.

In summary, we identify *de novo* enhancer activation and AP-1 as novel and key elements that pioneer and shape a transcriptionally permissive enhancer landscape in senescence.

### AP-1 pioneer TF bookmarking of senescence enhancer landscape foreshadows the senescence transcriptional program

Given our unexpected finding that most of the enhancer activation occurred *de novo* out of unmarked chromatin territories, *i.e.*, devoid of enhancer-related histone modifications H3K4me1 and H3K27ac and ending in an active H3K4me1^+^/ H3K27ac^+^ enhancer state at 144h, and that AP-1 TFs act as pioneers to shape the senescence enhancer landscape, we explored a possible role of AP-1 as a general bookmarking agent for future and past enhancer activity. Quantification of enhancer mark dynamics (Figure 3A and Supplementary Figures 3A-C) unveiled that for windows shifting from the “unmarked” state at T_0_ to an “active enhancer” state (H3K4me1^+^ / H3K27ac^+^) at either 72 h or 144 h, i.e. “*de novo* enhancers”, there is both a gradual increase in H3K4me1 and H3K27ac levels from initial levels (T_0_) similar to steady-state unmarked regions but different from poised enhancers, to final levels (144 h) indistinguishable from constitutive enhancers (Figure 3A and Supplementary Figures 3A-B). By contrast, for windows shifting from an “active enhancer” state at T_0_ to an “unmarked” state at either 72 h or 144 h, that we refer to as “remnant enhancers”, there is a progressive decrease both in H3K4me1 and H3K27ac levels from initial levels indistinguishable from constitutive enhancers to final levels similar to unmarked regions and distinct from poised enhancers (Figure 3A and Supplementary Figures 3A and 3C). The dynamic behavior of each enhancer class on average associated with the expression profile of nearby genes, with constitutive enhancers displaying constant gene expression, *de novo* enhancers increasing and remnant enhancers decreasing gene expression (Supplementary Figure 3D). To directly show the functional role of *de novo* enhancers we used a CRISPR interference (CRISPRi) approach^32,33^. Expression of 4 different gRNA targeting the dCas9-KRAB transcriptional repressor to *de novo* enhancers in the IL1α /IL1β genomic locus (g7, −14, −15, and −61) significantly reduced the expression of IL1β when analyzed 8 days after oncogenic RAS induction (Figure 3B). Interestingly, IL1α expression was only mildly reduced by the two gRNAs (g61 and g7) adjacent to the IL1β promoter (Figure 3B). While similar results were observed 14 days after oncogenic RAS induction (Supplementary Figure 3E), a control gRNA (g54) targeting a genomic region just downstream of the IL1α /IL1β locus did not affect either expression, while control gRNA guides g2 and g48, targeting sequences in-between two *de novo* enhancers, had only very moderate effects (Supplementary Figure 3F). Together, we render ample evidence that *de novo* and remnant enhancers are novel senescence-associated *cis*-regulatory modules that define the senescence transcriptional program.

**FIGURE 3:**
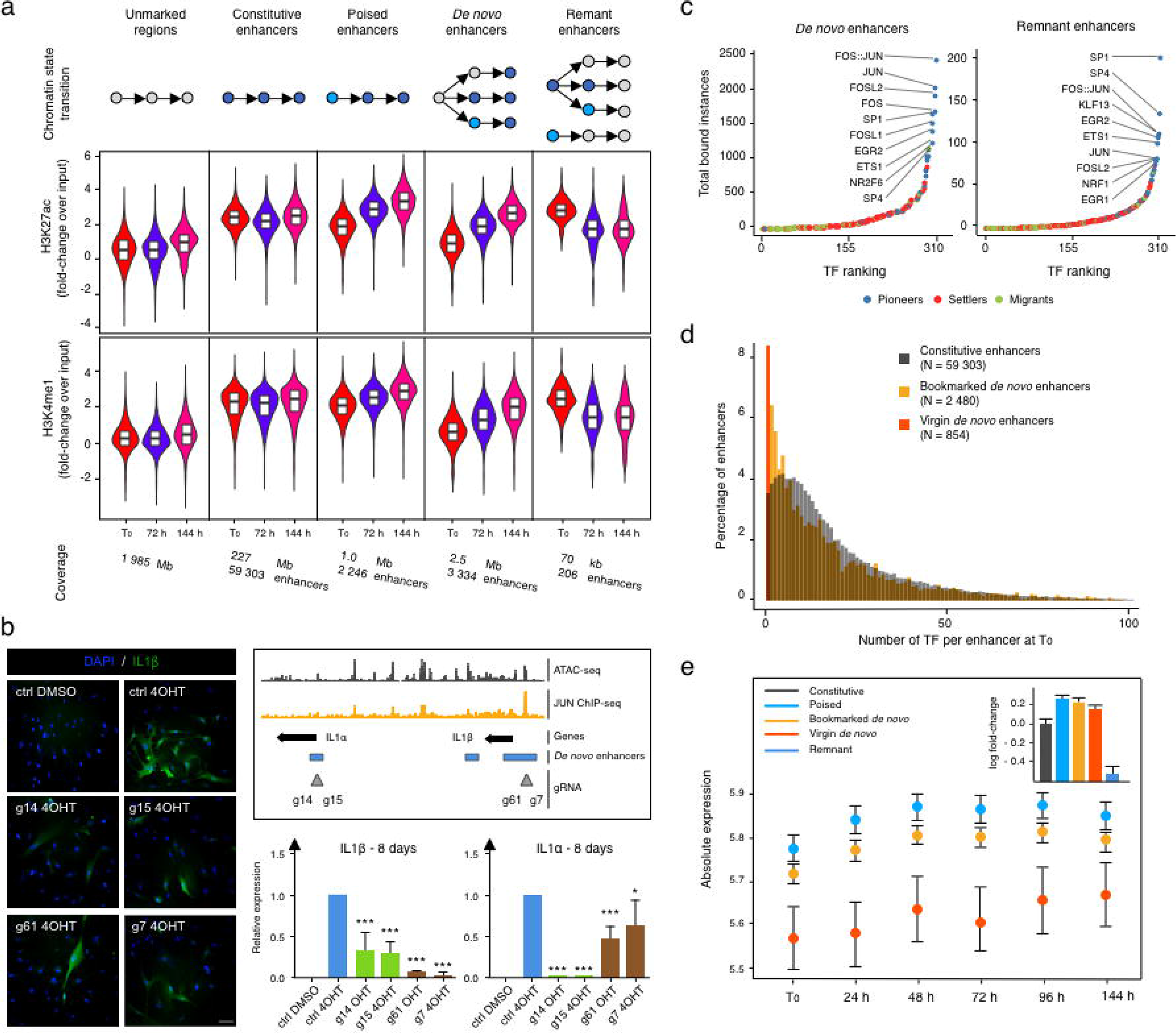
AP-1 pioneer TF bookmarking of senescence enhancer landscape foreshadows the senescence transcriptional program. **(a)** Distribution of fold-change in normalized enhancer marks H3K27ac and H3K4me1 ChIP-seq signals over input in the “unmarked”-, “constitutive”-, “poised”-, “*de novo”-*, and “remnant enhancers”-flagged genomic bins at indicated time-points (see Material and Methods for details). The cartoon at the top illustrates the temporal rules used to flag genomic bins. Bottom specifies the genomic coverage in mega bases (Mb) for each category and the corresponding number of enhancers. **(b)** WI38-ER: RASV12 were super-infected with dCas9-KRAB and individual guides (g14, 15, g61 and g7) targeting two *de novo* enhancers. Cells were pharmacologically selected and induced into RAS-OIS by 4-OHT. 8 days after RAS-OIS induction cells were stained by indirect immunofluorescence for IL1β or analyzed by RT-qPCR for the expression of IL1α or IL1β. WI38-ER: RASV12 treated with 4-OHT or DMSO served as positive and negative controls. Data represent mean ± SD (n=3). *p<0.05, ***p<0.001. Comparison with ctrl 4-OHT, one-way ANOVA (Dunnett’s test). Scale bar, 100 µm. **(c)** Rank plot depicting the summed occurrences for TF binding in *de novo* enhancers before RAS-OIS induction (left) and remnant enhancers after RAS-OIS (6 days) induction (right). Top ten TFs are indicated. **(d)** Distribution of total number (N) of TFs bound per enhancer for constitutive enhancers (grey), TF pre-marked *de novo* enhancers (yellow) and TF virgin *de novo* enhancers (orange). **(e)** Average absolute expression level (log_2_ scale) kinetics for genes associated with: poised (blue), TF pre-marked *de* novo (yellow), and TF virgin *de novo* enhancers (orange). Dots depict the average absolute expression level, and bars depict the standard error of the mean. Inset histogram illustrates the average leading log_2_ fold-change in expression (+/− standard error of the mean) for genes associated with constitutive (black), poised (light blue), TF pre-marked *de novo* (yellow) and TF virgin *de novo* (orange), and remnant enhancers (dark blue).

We next determined whether TFs bookmark *de novo* enhancers for future activation and also, whether TFs bookmark remnant enhancers after their inactivation as part of a molecular memory. Indeed, as shown in Figure 3C, we found that AP-1 is the predominant TF bookmarking *de novo* and remnant enhancers. Importantly, and highlighting the importance of AP-1 in bookmarking *de novo* enhancers for future activation, gRNAs chosen for CRISPRi were either overlapping with AP-1 binding sites (g14, g15 and g61) or in close proximity (g7), *i.e.* ~125bp outside of it (Figure 3B). Because CRISPRi can control repression over a length of two nucleosomes (~300bp)^34^, it is highly probable that g7 also affects this AP-1 binding site. Moreover, a control gRNA (g2) targeting a non-enhancer AP-1 site (Supplementary Figure 3F) did not affect IL1 expression strongly suggesting that only enhancer-positioned AP-1 sites are functional. Finally, we validated the importance of AP-1 TFs for *de novo* and remnant enhancer bookmarking by examining their positioning also in cells undergoing replicative senescence, which demonstrated that AP-1 TFs here also play a leading role for bookmarking (Supplementary Figure 3G). We conclude that AP-1 bookmarking of *de novo* and remnant enhancers is independent of oncogenic RAS signaling and a novel and cardinal feature that reflects future and past transcriptional activities in senescence.

While performing this analysis, we noticed that only 2,480 out of 3,334 *de novo* enhancers were TF bookmarked, while the remainder (n=854) lacked any detectable TF binding activity (Figure 3D). Thus, *de novo* enhancers can be further divided into two subclasses: 1) “TF bookmarked *de novo* enhancers” and 2) “TF virgin *de novo* enhancers” that are reminiscent to previously described latent enhancers^15,35^ expanding the senescence enhancer landscape. Next, we considered the chromatin state environment of the two *de novo* enhancer classes to further characterize them (Supplementary Figure 3H). While a chromatin state environment already rich in constitutive enhancers surrounded bookmarked *de novo* enhancers at T_0_ (*i.e.*, pre-OIS stimulation; left top and bottom plots), a chromatin state environment poor in constitutive enhancer elements surrounded virgin *de novo* enhancers at T_0_ (right top and bottom plots). Both AP-1 bookmarked and virgin *de novo* enhancers became progressively activated and expanded upon RAS-OIS induction. Given that AP-1 premarked *de novo* enhancers operate within pre-existing, active enhancer-rich *cis*-regulatory regions and virgin *de novo* enhancers in poor ones, we hypothesized that this might impact absolute gene expression levels and kinetics upon enhancer activation. Indeed, we observed that the nearest genes associated with bookmarked *de novo* enhancers were already expressed at higher basal levels (as were genes proximal to poised enhancers) and reached significantly higher absolute expression levels with faster kinetics after RAS-OIS induction. In contrast, virgin *de novo* enhancers showed only low-to-background basal expression levels and reached comparatively lower absolute expression levels with slower kinetics after RAS-OIS induction (Figure 3E). These results argue that TF bookmarking of *de novo* enhancers, similar to traditional enhancer poising^36^, is a chromatin-priming event that impacts gene expression kinetics and absolute output. Contrary to latent enhancers, our newly identified virgin enhancers do not serve an adaptive role in mediating stronger and faster gene expression upon restimulation as observed in macrophages^15^, but instead serve as novel enhancer elements for *de novo* gene expression. Finally, we plotted leading gene expression fold-changes against the number of *de novo* enhancers in a given prospective senescence enhancer region. Remarkably, we discovered that a single *de novo* enhancer element of 100 bp can substantially activate the expression of its nearest gene and that there exists a positive correlation between the number of *de novo* enhancer elements and the expression increase of their nearest genes (Supplementary Figure 3I).

Altogether, our results provide compelling evidence that *de novo* and remnant enhancers play a critical role for ensuring that genes pertinent for senescence are expressed at the correct time and the correct level and highlight the importance of AP-1 bookmarking for epigenetic memorization of past and future enhancer activity to define the senescence transcriptional program.

### A hierarchical TF network defines the senescence transcriptional program

The combinatorial and dynamic binding of TFs to enhancers and their organization into TF networks are central to the spatiotemporal specificity of gene expression and a key determinant in cell fate decisions^37^. TF networks are frequently corrupted in disease and thus, a detailed understanding on TF networks has important implications for developing and improving new therapeutic strategies^38^. Currently, a TF network regulating senescence is not available, which precludes a deliberate therapeutic manipulation of the senescence phenotype. Importantly, TF networks deduced *in silico* from the integration of time-resolved multidimensional, genome-wide datasets improve the accuracy and predictive power of such networks.

To elucidate the combinatorial and dynamic binding of TFs to enhancers and their organization into TF networks, we first computed co-occurring pairs of TFs in enhancers (Figure 4A, Supplementary Figure 4A and Supplementary data: see under Code availability in Material and Methods) followed by a topic machine learning approach that dissects the complexity of combinatorial binding of many TFs into compact and easily interpretable regulatory modules or TF “lexicons” that form the thematic structures driving the RAS-OIS gene expression program (Figure 4B)^39,40^. These analyses illustrated two key points. First, as shown in the co-binding matrix of Figure 4A and heatmap of Figure 4B, AP-1 pioneer TFs interact genome-wide with most of the remaining non-pioneer TFs (*i.e.*, settlers and migrant TFs; Figure 4A), have the highest total number of binding sites (Figure 4B, grey colored box plot) and contribute to virtually all of the 54 TF lexicons (Figure 4B, green colored boxplot) with lexicon 22 being the most frequently represented lexicon genome-wide (Figure 4B, top orange colored box plot). Our interactive heatmap of Figure 4B (Supplementary data: see under Code availability in Material and Methods) provides a valuable resource for generating new hypotheses to functionally dissect TF interactions in cells undergoing RAS-OIS. Second, TF lexicon usage associates with specific chromatin states (Supplementary Figure 4B). For example, lexicons 21 and 22 are exclusively used for enhancers holding most of the AP-1 binding instances, while lexicon 50 is strongly related to polycomb repressor complex (PRC)-repressed regions and lexicons 44 and 52 predominantly associate with promoters. Interestingly among the most prominent TFs in lexicon 50 are the known PRC-interacting transcriptional co-repressor complex REST and insulator CTCF^41,42^. The latter implies that these proteins may recruit PRC to silence or structure genomic regions, an intriguing possibility that deserves further investigation. Moreover, the promoter-centric lexicon 52 holds many E2F TFs, which is in line with a primary role of E2Fs at promoters^43^.

**FIGURE 4:**
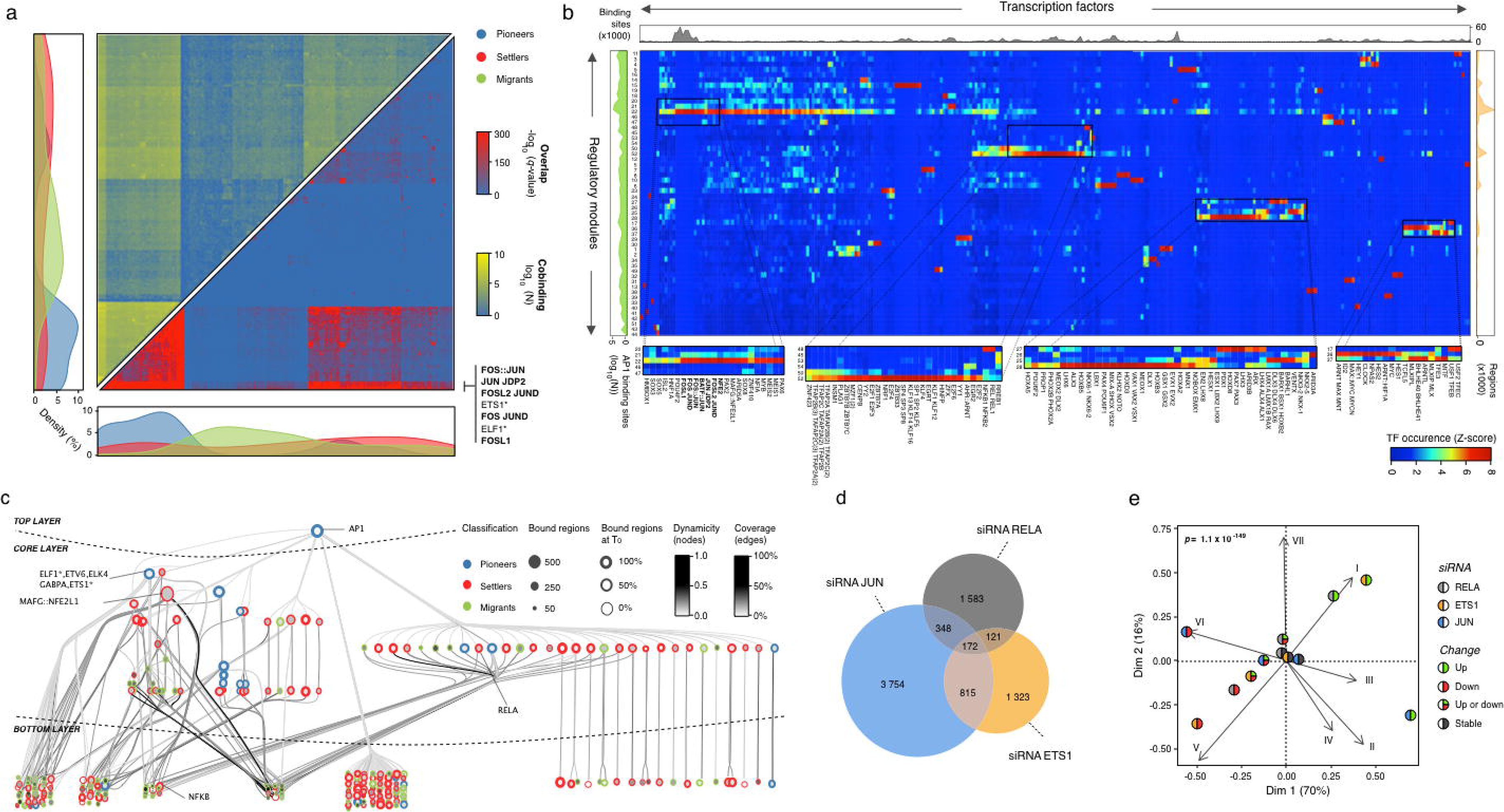
A hierarchical TF network defines the senescence transcriptional program. **(a)** Genome-wide transcription factor co-binding occurrence matrix summed across all time-points (left, shades from blue to yellow, in log_10_ scale). Overlap significance was calculated by a hyper-geometric test (right, shades from blue to red, in −log_10_ scale). The co-binding occurrence matrix was clustered using Ward’s aggregation criterion and corresponding, corrected *q*-values were projected on this clustering. The graphs on the left and bottom show the density in pioneer, migrant and settler TFs along each axis of the matrix. **(b)** Heatmap describing the association between individual TFs (row) and TF lexicons (columns). Four boxed out insets provide detailed information on TF composition of lexicons. A comprehensive, high-resolution and interactive heatmap is shown in Supplementary Data (see under Code availability in Material and Methods). The right bar plot shows the total number of binding sites for each TF. The top bar plot shows the total number of regions for each regulatory module. The bottom bar plot shows the average proportion of AP1 binding sites inside each regulatory module. **(c)** Graphical representation of the hierarchical TF network for transcriptomic module VI. Nodes (circles) represent TFs and an oriented edge (line) connecting TFs A and B means that at least 30 % of the regions bound by B were also bound by A at the same time point or before. In order to simplify the visualization, we represent strongly connected components (SCCs) as a single node and performed a transitive reduction (TR). Node color is based on the average dynamicity of the SCC members. Node border color indicates their classification as pioneer (blue), settler (red) or migrant (green). Node border thickness encodes the percentage of bound regions before RAS stimulation. Edge color was calculated accordingly to the relative coverage of the outgoing TF over the incoming TF. The network has three layers: top, core and bottom. Nodes in the top have no incoming edges and nodes in the bottom have no outgoing edges. The core layer comprises TFs that have both incoming and outgoing edges. Interactive Cytoscape graphs are accessible as Supplementary data (see under Code availability in Material and Methods). **(d)** Venn diagram showing the specificities and overlaps in differentially expressed direct target genes upon siRNA-mediated AP-1-c*JUN, ETS1,* and *RELA* depletion in RAS-OIS cells at day 6 (fully senescent cells). Genes are considered as direct targets of a given TF when PIQ predicts that the TF bound to an enhancer associates to this gene (see Materials and Methods for details). Promoters were excluded from the analysis. **(e)** Asymmetric biplot for correspondence analysis between transcriptomic clusters and the number of up-, down-, up-or-down- or nonregulated (stable) genes upon siRNA-mediated AP-1-c*JUN*, *ETS1* or *RELA* depletion. The *p-*value reflects the strength of the association as assessed with a χ^2^ test.

Next, we developed an algorithm, based on our temporal TF co-binding information and a previously published TF network (Supplementary Figure 4C)^44^, to visualize the hierarchical structure of the senescence TF network. In Figure 4C we show a representative example of the TF network of SASP gene module VI. The network has a three-layered architecture: i) a top layer defined exclusively by the AP-1 family of pioneer TFs ii) a core layer composed mostly of other pioneer and settler TFs, and iii) a bottom layer characterized by settler and migrant TFs (Figures 4C and Cytoscape interactive maps in Supplementary Data: see under Code availability in Material and Methods). The core layer itself separates into a multi-level and single-level core, depending on the complexity of TF connectivity to the top and bottom layers (Figure 4C). Remarkably, the organizational logic of the TF network is highly similar, if not identical, for all gene expression modules despite high transcription factor diversity in the core and bottom layers (Supplementary data: see under Code availability in Material and Methods). The TF network topology for RAS-OIS is congruent with the biochemical and dynamic properties of each TF category (*i.e.*, pioneer, settler or migrant) in each layer of the network. As the interactions flow from the top to the bottom layer, there is an increasing dynamicity and number of TFs and a decreasing number of bound regions (Supplementary Figures 4D-E). Ranking the dynamicity index and the number of bound regions for all TFs in each network confirmed the hierarchical principles of their organization, with a common core of highly connected TFs from the top and core layers shared across all networks (Supplementary Figure 4F, black circle at center). Variability in the composition of the most dynamic TFs of the core and bottom layers defines the gene expression module specificity for each network and its corresponding specialized transcriptional output (Supplementary Figures 4G-I). Thus, TF network topology imposes and constrains the position of a given TF in the network and thus, its gene-regulatory contribution. Our data also revealed unanticipated plasticity in transcription factor binding leading to similar gene expression, thus, refuting the simple rule that co-expression behooves co-regulation^45^.

Our hierarchical TF network model for RAS-OIS enhancers predicted that the number of direct target genes regulated by a given TF is a function of its position in the TF network hierarchy. To test this prediction, we performed transient RNA interference (siRNA) experiments targeting AP-1-cJUN (top layer), ETS1 (multi-level core layer) and RELA (single-level core layer) in fully senescent RAS-OIS cells (144 h), determined the global transcriptome profiles and compared them to the transcriptomes of cells transfected with a non-targeting siRNA (siCTRL) (Figure 4D). Consistent with the TF network hierarchy, silencing of AP-1-cJUN affected the most substantial number direct gene targets (n=5,089), followed by ETS1 (n=2,431) and RELA (n=2,224), thus, confirming the master regulatory role of AP-1 pioneer TFs at enhancers and in the execution of the RAS-OIS gene expression program. Specifically, 172 genes were co-regulated by the three TFs, while 987 were co-regulated by cJUN and ETS1, 520 by JUN and RELA, and 293 by ETS and RELA. Correspondence analysis (CA) revealed that perturbing the function of AP-1-cJUN, ETS1 or RELA could separate faithfully (p = 1.8 × 10^−149^) up-regulated (V-VII) from down-regulated gene expression modules (I-IV) (Figure 4E), which aligns perfectly, both with the CA for chromatin states (see Figures 2A-B) and the differential impact of the TFs on RAS-OIS-associated enhancer activation as predicted in the TF network analysis (Figure 4C and Supplementary data: see under Code availability in Material and Methods).

We conclude that the senescence response is encoded by a universal three-layered TF network architecture and relies strongly on the exploitation of an enhancer landscape implemented by AP-1 pioneer TFs to choreograph the OIS transcriptional program *via* local, diverse and dynamic interactions with settler and migrant TFs.

### Hierarchy Matters: Functional Perturbation of AP-1 pioneer TF, but no other TF, reverts the senescence clock

Pioneer TFs have been identified as important drivers of cell fate changes during adaptive and cellular reprogramming as well as in cells undergoing malignant transformation^46,47^. As such, they represent attractive targets to manipulate cell fate for diverse research and therapeutic purposes^19^.

The identification of AP-1-cJUN as a principal pioneer TF in fibroblasts undergoing RAS-OIS raised the possibility that perturbing its function could considerably change the transcriptional trajectory of the OIS cell fate, while perturbation of other TFs should not. To test this hypothesis, we depleted AP-1-cJUN, ETS1 and RELA at T_0_, 72 h and 144 h following oncogenic RAS expression and compared global gene expression profiles with siCTRL treated cells at identical time-points. Capturing their transcriptional trajectories using PCA illustrated that functional perturbation of ETS1 and RELA shifted trajectories along the second principal component (PC2, which captures siRNA-related variability) at any given time-point compared to the control time course, but it did not affect the timely execution of the RAS-OIS gene expression program, since there is not shift along the first principal component (PC1, which captures time-related variability). By contrast, perturbing AP-1-cJUN function shifted trajectories both along PC1 and PC2 and effectively reverted the RAS-OIS transcriptional trajectory to a profile closely related to that of siCTRL-treated fibroblasts at 72 h after RAS-OIS induction. Silencing AP-1-cJUN expression at 72 h also pushed the transcriptional profile closer to control-treated cells at day T0 (Figure 5A, blue arrow). Functional overrepresentation analyses of the target genes (direct and/or indirect) of each TF further supported the siJUN-mediated reversion of the RAS-OIS transcriptional trajectory demonstrating that depletion of AP-1-cJUN strongly affected both the repression of the inflammatory response (*i.e.*, the SASP) and a partial reactivation of pro-proliferation genes (*i.e.,* E2F, G2M and mitotic spindle targets) (Figure 5B and Supplementary Figures 5A-C). A complete exit of senescence is not expected here, however, as AP-1 is absolutely required for proliferation^48,49^.

**FIGURE 5:**
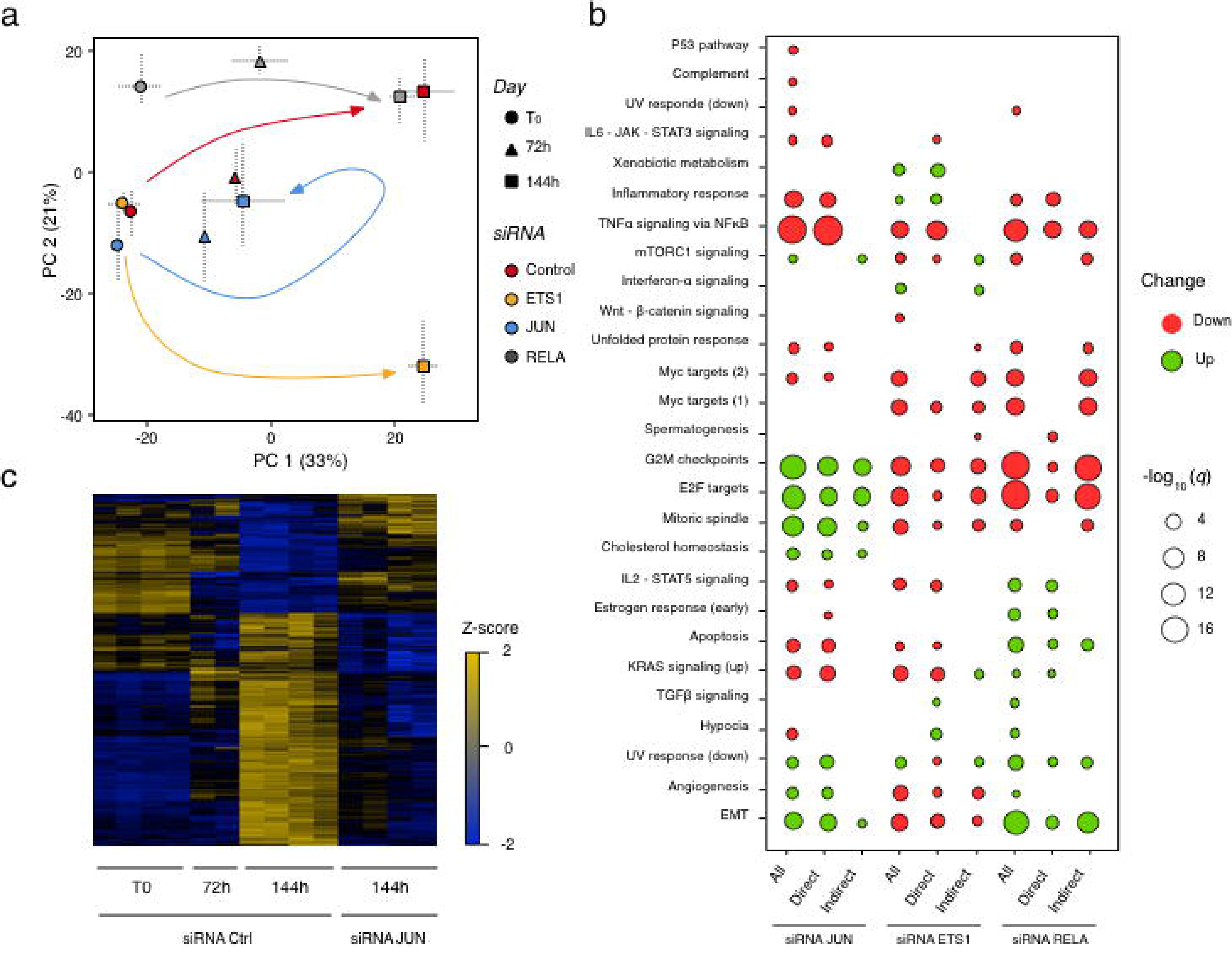
Hierarchy Matters: Functional Perturbation of AP-1 pioneer TF, but no other TF, reverts the senescence clock. **(a)** Principal component analysis (PCA) on transcriptomes obtained from siRNA-mediated depletion of AP-1-c*JUN*, *ETS1* or *RELA* at indicated timepoints of the RAS-OIS timecourse. Horizontal and vertical bars show minimal and maximal coordinates for each siRNA and time-point on principal components one (PC1, horizontal axis) and two (PC2, vertical axis). **(b)** Functional overrepresentation map showing Molecular Signature Database (MSigDB) hallmark pathways associated to “All, Direct Target and Indirect Target” genes differentially expressed after siRNA-mediated AP-1-c*JUN*, *ETS1* or *RELA* depletion. Genes are considered as direct targets when a PIQ prediction for the given TF is falling inside an enhancer associated to this specific gene. Promoters are excluded from the analysis. The size of dots is proportional to the −log_10_ *q*-value based on the hypergeometric distribution obtained when testing for over-representation, and their color denote whether the term is enriched for an up or down-regulated gene list. **(c)** Heatmap comparing gene expression profiles of siRNA-Control-treated (siCTRL) cells at indicated time-points of OIS and siRNA-cJUN treated senescent RAS-OIS cells at day 6 (144h).

To quantify and visualize the temporal overlaps in differentially expressed genes between siJUN and siCTRL-treated cells we used an UpSet plot (Supplementary Figures 5D) and expression heatmaps (Figure 5C and Supplementary Figures 5E-G). Congruent with a resetting of the senescence clock, a significant number of pro-proliferation E2F target genes (14%; *e.g. CCNB2* or *CDCA8*) were up-regulated (Supplementary Figure 5E) and NF-κB-regulated SASP target genes (*e.g. IL1B* or *IL6*) were down-regulated (60%) (Supplementary Figure 5F). cJUN-depleted RAS-OIS cells also shared a similar expression profile for a subset of genes (27%) of the Notch-1 intracellular domain (NC1ID)-induced senescence (NIS) transcriptional signature^50^ that develops within the first 72-96 h of RAS-OIS (Supplementary Figure 5G). Thus, AP-1 inhibition is a powerful and save means to potently repress SASP expression in senescent cells without affecting their cell cycle arrest.

Altogether, these data identify AP-1 as a master regulator and molecular “time-keeper” of senescence. Our detailed description of the layered TF network architecture will facilitate targeted disruption of TFs to manipulate specific features of the senescence phenotype for future therapeutic benefit.

## DISCUSSION

Exploiting senescence targeting for treating age-related diseases and cancer requires a detailed knowledge of the transcriptional, epigenetic, and signaling mechanisms defining the basis and realization of the senescence program, which is currently missing. To fill this critical gap in our knowledge, we used a dynamic, multidimensional approach at high-resolution to define the gene-regulatory code driving the senescence cell fate.

A central finding of our study is that the senescence program is defined and driven by a predetermined enhancer landscape that is sequentially (in)activated during the senescence process. AP1 is instrumental for this predetermination by imprinting a prospective senescence enhancer landscape that, in the absence of traditional enhancer histone-modification marks, foreshadows future transcriptional activation. This is a surprising discovery given that AP-1 transcriptional activation has been traditionally linked to growth-factor and MAPK signaling^51^. There is, however, now accumulating evidence that AP-1 also plays an essential role as a pioneering factor for establishing cell type-specific enhancers and cellular identities^52,53^. In line with its role in pioneering and bookmarking enhancers, we show that AP-1 is also recruited *de novo* as a first line TF to “virgin” enhancers and that it serves as a molecular memory for enhancers that become inactivated during the senescence fate transition that we termed “remnant” enhancers. Based on these findings we propose a model by which “enhancer recycling” of AP1-bookmarked future and past enhancer activities is an evolutionary conserved mechanism that allows for modular and flexible, yet, efficient transcriptional responses to incoming signals. We stipulate that the senescence program is preserved through AP-1 binding to enhancer chromatin as part of epigenetic memory of the cell’s developmental (stress) history bypassing histone modification-dependent bookmarking to store genomic information. Further, given the pristine specificity of the newly identified prospective and remnant enhancers they can be used as urgently needed specific, rather than associated, senescence biomarkers and to predict a cell’s potential to undergo senescence. This latter carries also important translational implications for identifying cancers that would respond positively to pro-senescence therapy. A natural question that arises from our data is whether the senescence program is universal to all inducing stimuli and cell types or if multiple senescence programs exist. Based on the data presented here and work in progress, we predict that the organizational principles of the senescence program we defined here hold for all cell types and inducers. Additional time-resolved studies of various inducers in different cell types are required and currently ongoing to answer this question definitively.

Another key finding is the reversibility of senescence by an informed intervention on network topology that we delineated in this study. Indeed, silencing the function of a single TF sitting atop the TF network hierarchy, AP-1, is sufficient to revert the “senescence clock”. We thus define after the “telomere clock” a second, “epigenomic-enhancer clock” regulating the senescence process. Why does functional AP-1 perturbation not lead to complete senescence exit? Based on published^49^ and our own results we surmise that AP1 depletion does not lead to a full cell cycle re-entry and proliferation, because AP1 plays important roles for proliferation. Thus, AP1 confines cells to their existing proliferative state and therefore may be viewed as a ‘locking device’ that restricts cells to their current state. However, we provide compelling evidence that functional inhibition of AP1 factors reverts the senescence transcriptional program and potently represses the expression of the pro-inflammatory senescence-associated secretory phenotype (SASP). This finding has great therapeutic potential, as pharmacological interference with AP1 using selective inhibitors (*e.g.* improved T-5224 derivatives) would allow to control effectively the detrimental effects of the SASP in promoting cancer and other age-related diseases^54–56^. In summary, we believe that AP1 is a prime target for therapeutic SASP modulation *in vivo*.

By determining the layered architecture/organizational principles of the TF network that orchestrate(s) the transition to OIS, we revealed the plasticity and stability of the senescent phenotype. We show that a highly flexible, combinatorial TF interactome establishes the senescence program, which is in line with the TF network dynamics during hematopoietic and stem cell differentiation^57,58^. In addition, we demonstrate that targeted engineering of specific nodes at different layers of the TF network disrupts gene expression with a corresponding magnitude, suggesting a path for the manipulation of the senescent phenotype *in vivo*. Pharmacological inhibition of TFs (see above for AP-1), signal transduction molecules, such as kinases or acetylases that converge in the activation of TFs, could represent a viable approach for manipulating the senescent phenotype *in vivo*^59^. Alternatively, small molecules that prevent TF-TF combinatorial interactions could also be envisioned^60^.

In conclusion, the present work emphasizes the advantages of, and indeed the need for, integrating time-resolved genome-wide profiles to describe and interrogate the transition to senescence. This approach generates detailed knowledge necessary to develop strategies for manipulating/engineering the senescent cell fate (and other cell fate transitions) *in vivo* for research and therapeutic purposes. Overall, our study provides a comprehensive resource for the generation of novel hypotheses regarding senescence regulation, offers important mechanistic, regulatory insights that could translate to the study of other cell fate transitions and provide new inroads for the diagnosis and manipulation of the senescence state in age-related diseases and cancer.

## MATERIAL AND METHODS

### Cell culture

WI-38 fibroblasts (purchased from ECCAC) were cultured in Dulbecco’s Modified Eagle’s medium (DMEM) containing 10% FBS and 1X Primocin (Invivogen) at 37°C and 3% oxygen. WI-38-ER: RASV12 fibroblasts were generated by retroviral transduction as previously described^9^. Senescence was induced by addition of 400 nM 4-hydroxytamoxifen (4-OHT, Sigma Cat no. H7904-5MG) to the culture medium and samples were collected and processed at the time points indicated in the main text. Replicative senescent cells were generated by proliferative exhaustion and were used for experiments when cell cultures went through 1 population (PD) per 3 weeks, were >80% positive for senescence-associated beta galactosidase activity (SABG) and stained negative for EdU (see below for details). For the induction of quiescence, WI-38 fibroblasts were cultured in DMEM containing 0.2% FBS for up to 4 consecutive days and samples were collected and processed as described in the main text.

### ATAC-seq

The transposition reaction and library construction were performed as previously described^24^. Briefly, 50,000 cells from each time point of the senescence time course (2 biological replicates) were collected, washed in 1X in PBS and centrifuged at 500 × *g* at 4°C for 5 min. Nuclei were extracted by incubation of cells in Nuclear Extraction Buffer (NEB) containing 10 mM Tris-HCl, pH 7.4, 10 mM NaCl, 3 mM MgCl2, 0.1% IGEPAL CA-630 and immediately centrifuging at 500 × *g* at 4°C for 5 min. The supernatant was carefully removed by pipetting, and the transposition was performed by resuspension of nuclei in 50 µL of Transposition Mix containing 1X TD Buffer (Illumina) and 2.5 µL Tn5 (Illumina) for 30 min at 37°C. DNA was extracted using the QIAGEN MinElute kit. Libraries were produced by PCR amplification (12-14 cycles) of tagmented DNA using the NEB Next High-Fidelity 2x PCR Master Mix (New England Biolabs). Library quality was assessed in an Agilent Bioanalyzer 2100. Paired-end sequencing was performed in an Illumina Hiseq 2500. Typically, 30-50 million reads per library are required for downstream analyses.

### Histone modification and transcription factor ChIP-seq

WI-38-ER: RASV12 fibroblasts were treated with 400 nM 4-OHT for 0, 72 and 144 hours, and 10^7^ cells (per time point, minimum two biological replicates) were fixed in 1% formaldehyde for 15 min, quenched in 2M glycine for additional 5 min and pelleted by centrifugation at 2,000 rpm, 4°C for 4 min. For histone modification ChIP-seq, nuclei were extracted in Extraction Buffer 2 (0.25 M sucrose, 10 mM Tris-HCl pH 8.0, 10 mM MgCl_2_, 1% Triton X-100 and proteinase inhibitor cocktail) on ice for 10 min followed by centrifugation at 3,000 × *g* at 4°C for 10 min. The supernatant was removed and nuclei were resuspended in Nuclei Lysis Buffer (50 mM Tris-HCl pH 8.0, 10 mM EDTA, 1% SDS and proteinase inhibitor cocktail). Sonication was performed using a Diagenode Picoruptor until the desired average fragment size (100-500 bp) was obtained. Soluble chromatin was obtained by centrifugation at 11,500 rpm for 10 min at 4°C and chromatin was diluted 10-fold. Immunoprecipitation was performed overnight at 4°C with rotation using 1-2 × 10^6^ cell equivalents per immunoprecipitation using antibodies (5 µg) against H3K4me1 (Abcam), H3K27ac (Abcam), H3K4me3 (Millipore), H3K27me3 (Millipore). Subsequently, 30 µL of Ultralink Resin (Thermo Fisher Scientific) was added and allowed to tumble for 4h at 4°C. The resin was pelleted by centrifugation and washed three times in low salt buffer (150 mM NaCl, 0.1% SDS, 1% Triton X-100, 20 mM EDTA, 20 mM Tris-HCl pH 8.0), one time in high salt buffer (500 mM NaCl, 0.1% SDS, 1% Triton X-100, 20 mM EDTA, 20 mM Tris-HCl pH 8.0), two times in lithium chloride buffer (250 mM LiCl, 1% IGEPAL CA-630, 15 sodium deoxycholate, 1 mM EDTA, 10 mM Tris-HCl pH 8.0) and two times in TE buffer (10 mM Tris-HCl, 1 mM EDTA). For transcription factor ChIP-seq, fibroblasts were treated as described above except that chromatin was isolated using the enzymatic SimpleChIP kit (Cell Signaling) according to the manufacturer’s instructions, obtaining chromatin with an average fragment length of 4-5 nucleosomes. Immunoprecipitation was performed overnight at 4°C with rotation using 6-10 × 10^6^ cell equivalents per immunoprecipitation using antibodies (5 µg) against cJUN, FOSL2 and RELA (Santa Cruz Biotechnologies) and processed as described above. Washed beads were resuspended in elution buffer (10 mM Tris-Cl pH 8.0, 5 mM EDTA, 300 mM NaCl, 0.5% SDS) treated with RNAse H (30 min, 37 °C) and Proteinase K (2 h, 37°C), 1 µL glycogen (20 mg/mL, Ambion) was added, and decrosslinked overnight at 65 °C. For histone modifications, DNA was recovered by mixing the decrosslinked supernatant with 2.2X SPRI beads followed by 4 min incubation at RT. The SPRI beads were washed twice in 80% ethanol, allowed to dry, and DNA was eluted by in 35 µL 10 mM Tris-Cl pH 8.0. For transcription factors, DNA was eluted by phenol/chloroform extraction (2X) followed by ethanol precipitation overnight at −20°C. The DNA pellet was washed with 70% ethanol, allowed to dry, and DNA was resuspended in 35 µL 10 mM Tris-Cl pH 8.0. Histone modification libraries were constructed using the NextFlex ChIP-seq kit (Bioo Scientific) according to the manufacturer’s instructions. Libraries were amplified for 12 cycles. Transcription factor libraries were constructed using a modified protocol from the Accel-NGS 2S Plus DNA Library Kit (#21024), where we performed DNA extraction at each step using 25:24:1 phenol:chloroform:isoamyl alcohol followed by overnight ethanol precipitation of DNA at each step of the protocol. Additionally, we enriched for small DNA fragments using AMPure-XP beads (Beckman-Coulter (#A63881). Libraries were then resuspended in 20 µL of low EDTA-TE buffer. Libraries were quality controlled in an Agilent Technologies 4200 Tapestation (G2991-90001) and quantified using the Invitrogen Qubit DS DNA HS Assay kit (Q32854). Libraries were sequenced using an Illumina High-Seq 2500. Typically, 30-50 million reads were required for downstream analyses.

### RNA and microarrays

RNA from each time point from the senescence and quiescence time series (2 biological replicates) was purified using the QIAGEN RNeasy Plus kit according to the manufacturer’s instructions. 100 ng RNA per sample was analyzed using Affymetrix Human Transcriptome Arrays 2.0, according to the manufacturer’s instructions.

### EdU staining and senescence-associated beta galactosidase activity (SABG)

Representative samples from the senescent and quiescent time series were evaluated for EdU incorporation using the Click-iT EdU Alexa Fluor Imaging Kit (Thermo Fisher Scientific) according to the manufacturer’s instructions. SABG activity was assessed as previously described^61^. Cells were imaged in a Zeiss confocal fluorescence microscope and images analyzed using the ZEN suite.

### RNA interference

Small interfering RNAs (20 µM) targeting JUN (QIAGEN, Ambion), ETS1 (QIAGEN) and RELA (QIAGEN) as well as non-targeting controls were transfected into WI-38-ER: RASV12 using siIMPORTER reagent (Millipore) according to the manufacturer’s instructions (minimum 2 biological replicates per transcription factor per time course experiment). RAS-OIS was induced with 400 nM 4-OHT concomitantly with the addition of DMEM containing 20% FBS 4 hours after transfection and incubated overnight. Sixteen hours after transfection, cells were replenished with new media containing 10% FBS and 400 nM 4-OHT, and RNA was isolated at indicated time points and analyzed in Affymetrix Human Transcriptome Arrays 2.0.

### Expression microarray pre-processing

Raw Affymetrix HTA 2.0 array intensity data were analyzed using open-source Bioconductor packages on R. The quiescence and the RAS-OIS time series data were normalized together (2 conditions, 2 biological replicates per condition, 6 time points per replicates) using the robust multi-array average normalization approach implemented in the *oligo* package. Internal control probe sets were removed and average expression deciles over time-points were independently defined for each treatment. Probes whose average expression was lower than the 4^th^ expression decile in both conditions were removed for subsequent analyses. To remove sources of variation and account for batch effects, data were finally corrected with the *sva* package. To recover as much annotation information as possible, we combined Affymetrix HTA 2.0 annotations provided by Affymetrix and Ensembl through the packages *hta20sttranscriptcluster.db* and *biomaRt*. Principal component analysis and bi-clustering based on Pearson’s correlation and Ward’s aggregation criterion were used to confirm consistency between biological replicates and experimental conditions at each step of the pre-processing.

### Self-organizing maps (SOM)

Normalized log-scaled and filtered expression values were processed using the unsupervised machine learning method implemented in *oposSOM*^25^ to train a self-organizing map. This algorithm applies a neural network algorithm to project high dimensional data onto a two-dimensional visualization space. In this application, we used a two-dimensional grid of size 60 × 60 metagenes of rectangular topology. The SOM portraits were then plotted using a logarithmic fold-change scale. To define modules of co-expressed meta-genes, we used a clustering approach relying on distance matrix and implemented in *oposSOM*. Briefly, clusters of gene expression were determined based on the patterns of the distance map which visualizes the mean Euclidean distance of each SOM unit to its adjacent neighbors. This clustering algorithm referred to as D-clustering – finds the SOM units referring to local maxima of their mean distance with respect to their neighbors. These pixels form halos edging the relevant clusters in the respective distance map and enable robust determination of feature clusters in the SOM. We finally performed a gene set over-representation analysis in each cluster considering the Molecular Signature Database (MSigDB) hallmark gene sets using a right-tail modified Fisher’s exact test and the hypergeometric distribution to provide *p*-value.

### Correlation and multidimensional analyses

To highlight differences in expression profiles between quiescence and RAS-OIS through time, we used multi-dimensional scaling plot representing leading fold change, which is defined as the root-mean-square average of the log-fold-changes for the genes best distinguishing each pair of samples. To quantify the evolution of transcriptomic variability and noise through time, we looked at the gene expression density distributions for all possible pairs of treated *vs* T_0_ transcriptomes. Distributions were estimated using kernel density estimation of all genes’ expression in the *i*^th^ T_0_ transcriptome and the *j*^th^ treated transcriptome. We also computed Pearson’s correlation for each of these combinations. The Pearson’s correlation between two transcriptomes, *X* and *Y* containing *n* gene expressions, is obtained by 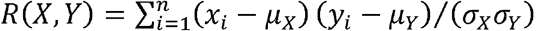, where *x*_i_ and *y*_i_ are the *i*^th^ observation in the vectors *X* and *Y* respectively, *µ_X_* and *µ_Y_* the average values of each transcriptome, and σ*_X_* and σ*_y_*, the corresponding standard deviations.

### Information theory – derived metrics

To evaluate transcriptome diversity and specialization, we used an approach based on information theory as described in ^26^.

### Gene expression time series analysis

Normalized log-scaled and filtered expression data related to the quiescence and the OIS time series were further considered for differential analysis with *limma*^62^. To define an RAS-OIS specific transcriptomic signature, we proceeded in three steps, each relying on linear mixed model cubic B-splines, as nonlinear response patterns are commonly encountered in time course biological data. For each probe, and each treatment the expression was modeled as follow:

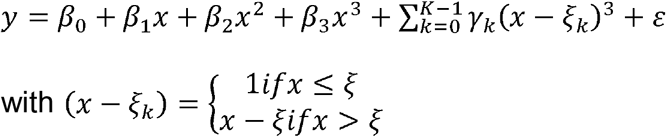

where *β*_0_ is the average probe expression over all samples in a given condition, *β*_1 − 3_ the model coefficients, *K* the number of knots, *ξ*_k_ the *k*^th^ knot and ε the error term. First, we defined probes responding over time to RASV12 induction. Second, we considered all together the quiescence and the RAS-OIS time series, as well as the interaction between time and treatment, and defined probes responding to one or the other treatment over time, as well as probes responding differently between the two treatments at any time point. We finally defined the set of probes responding consistently to both treatment and time and removed these probes from the global set of probes responding to RASV12 induction defined at the first step. Moderated *F*-statistics that combine the empirical Bayes moderated *t*-statistics for all contrasts into an overall test of significance for each probe were used to assess the significance of the observed expression changes. At any step of this workflow, *p*-values were corrected for multiple testing using the FDR approach for a stringent significance level of 0.005. For validation purposes, we also wanted to compress the RAS-OIS time-series and achieve a volcano plot representation. To do so, we’ve computed the maximal absolute log_2_ fold change in expression in the RAS-OIS time series considering T_0_ as the reference and selected up and down regulated probes using an absolute log_2_ fold change cutoff at 1.2 and a corrected *p*-value cutoff of 0.005. We then build a scatter-plot plotting the log_10_ significance versus log_2_ fold-change on the y and x axes, respectively. Probes responding consistently to both ER: RASV12 induction and quiescence were finally over-plotted.

### Gene expression unsupervised clustering

Probes constitutive of the RAS-OIS specific transcriptomic signature were clustered using the weighted gene correlated network analysis approach implemented in the *WGCNA* R package^63^. Standard WGCNA parameters were used for the analysis, with the exceptions of soft-thresholding power which was defined using methods described by and set at 18. The 7 co-expressed probe clusters identified were further functionally characterized using gene set over-representation tests. The same approach as previously described for the SOM-defined clusters was used.

### Histone modification ChIP-seq data processing

Reads were cleaned and trimmed using *fastq-mcf* from the *ea-utils* suite v1.1.2 to remove adapters, low quality bases and reads, and discard reads shorter than 25 bp after filtering. Reads were then aligned to the human reference genome (hg19) with *bowtie* v1.1.1 using best matches parameters (bowtie -v 2 -m 1 --best --strata). Alignment files were further processed with *samtools* v1.2 and *PicardTools* v1.130 to flag PCR and optical duplicates and remove alignments located in Encode blacklisted regions. Fragment size was estimated *in silico* for each library using *spp* v1.10.1. Genome-wide consistency between replicates was checked using custom R scripts. Enriched regions were identified for each replicate independently with *MACS* v2.1.0 with non-IPed genomic DNA as a control (macs2 callpeak --nomodel --shiftsize --shift-control --gsize hs -p 1e-1). These relaxed peak lists were then processed through the irreproducible discovery rate (IDR) pipeline^64^ to generate an optimal and reproducible set of peaks for each histone modification and each time point.

### ATAC-seq data processing

Paired-ends reads were cropped to 100bp with *trimmomatic* v0.36^65^ and cleaned using *cutadapt* v1.8.3^66^ to remove Nextera adapters, low quality bases and reads, and discard reads shorter than 25 bp after filtering. Fragments were then aligned to the human reference genome (hg19) using *bowtie2* v2.2.3 discarding inconsistent pairs and considering a maximum insert size of 2kb (bowtie2 -N 0 --no-mixed --no-discordant -- minins 30 --maxins 2000). Alignment files were further processed with *samtools* v1.2 and *PicardTools* v1.130 to flag PCR and optical duplicates and remove alignments located in Encode blacklisted regions. Accessible regions were identified using *MACS2* v2.1.0 without control (macs2 callpeak --gsize hs -p 1e-3). These relaxed peak lists were then processed through the irreproducible discovery rate (IDR) pipeline to generate an optimal and reproducible set of peaks for each time point.

### Normalized ATAC-seq and ChIP-seq signal tracks

After verifying the consistency between biological replicates, time points and data type using *deepTools*^67^, alignments related to biological replicates for a given assay and a given time point were combined. We then binned the genome in 200bp non-overlapping windows and generated genome-wide read count matrices for each assay independently. These matrices were finally quantile normalized with custom R script and further used to generate genome-wide signal tracts.

### Histone modification ChIP-seq and ATAC-seq differential analysis

After assessing library saturation using *preseqR*, alignment and peak data were imported and pre-processed in R using the *DiffBind* package^68^. Briefly, for a given histone modification type, we first defined the global reproducible peak set as the union of each time-specific reproducible peak sets defined previously. We then counted the number of reads mapping inside each of these intervals at each time point and for each replicate. The raw count matrix was then normalized for sequencing depth using a non-linear full quantile normalization as implemented in the *EDASeq* package^69^. To remove sources of unwanted variation and consider batch effects, data were finally corrected with the *RUVSeq*^70^ package considering 2 surrogate variables. Differential analyses for count data were performed using *edgeR*^71^ considering time and batch in the design matrix, by fitting a negative binomial generalized log-linear model to the read counts for each peak. Peaks were finally annotated using *ChIPpeakAnno* considering annotations provided by Ensembl v86.

### Chromatin state differential analysis

To quantify and define combinatorial chromatin state dynamics in space and time, we analyzed histone modification combinations with the *chromstaR* package^72^. Briefly, after partitioning the genome into 100bp non-overlapping bins and counting the number of reads mapping into each bin at each time point and for each histone modification, this algorithm relies on a univariate Hidden Markov Model (HMM) with two hidden states (unmodified, modified). This HMM is used to fit the parameters of the two-component mixture of zero-inflated negative binomial distribution considered to model read counts for every ChIP-seq experiments. A multivariate HMM is then used to assign every bin in the genome to one of the multivariate components considering 2^(3 time points × 4 histone modifications)^ possible states. To limit computational burden and focus on accurate differences, the analysis was run in differential mode with a 100bp resolution (*i.e.* smaller than a single nucleosome), such that every mark is first analyzed separately with all conditions combined while the full combinatorial state dynamics is rebuilt by combining the differential calls obtained for the four marks. We finally filtered out differential calls not overlapping with any histone modification and ATAC-seq reproducible peaks. To properly associate histone modification combinations with biologically meaningful mnemonics, we made an extensive comparison between the binning we obtained in WI38 fibroblasts undergoing RAS-OIS and IMR90 fetal lung fibroblasts chromatin states described in the scope of the Epigenomic Roadmap consortium. To test for association between changes in chromatin states through time and gene expression modules we ran a correspondence analysis. Briefly, genomic loci experiencing changes in chromatin states through time were first associated to the nearest gene. We then specifically focused on loci associated to genes belonging to any expression module and built a two-way contingency table summarizing the number of transition in states (considering all possible combinations) occurring in each expression module, further used as an input for a correspondence analysis using *FactoMineR*^73^. The significance of association between the two qualitative variables (transition in state and module) was assed using a *χ*^2^ test. Results of the CA were visualized using a row-metric-preserving contribution asymmetric biplot and filtering for the top contributing and well-projected (squared cosine > 0.5) changes in chromatin states.

### Motif enrichment analysis in active enhancers

For each time point independently, we defined the set of active enhancers as the overlap between H3K4me1, H3K27ac and ATAC-seq reproducible peaks using *bedtools*^74^. We then ran 3 independent motif enrichment analyses with *homer* v4.9^75^ using default parameters.

### Transcription factor footprinting

All transcription factor Position-Weight Matrices (PWM) representing eukaryote transcription factors were downloaded from the JASPAR database and used as an input for PIQ^21^ to predict transcription factor binding sites from the genome sequence on down-sampled ATAC-seq alignments. For each motif, we retained only binding sites that were within the reproducible ATAC-seq peaks and passed the default purity cut-off (70%). We then computed pairwise PWM similarities based on Pearson’s correlation, and clustered together PWMs sharing more than 90% similarity, defining a set of 310 non-redundant and distinct PWMs. The Pearson’s correlation between two PWM *P*^1^ and *P*^2^ of length *l* was defined as:

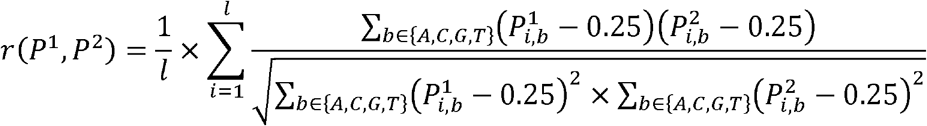

We further combined the bound instances identified with PIQ according to the PWM clustering.

### Transcription factor metrics

For each transcription factor, we computed the chromatin-opening index (COI), the motif dependence and the chromatin dependence (CD) following the approach described in^21^.

### Validation of PIQ predictions through ChIP-seq

To compare PIQ prediction with RELA, JUN and FOSL2 ChIP-seq data, we first used the approach suggested in ^21^, computing how many of the total ChIP-seq peaks are overlapping with any potential factor motif (since ChIP-Seq peaks can result from co-factor binding, and methods such as digital genomic footprinting are factor agnostic). We then used a more sophisticated approach aiming at correlating the ChIP-seq signal intensity with the bound / unbound status at PWM matches. For a given transcription factor (cJUN, FOSL2 or RELA,), we first considered all the PWM matches located inside ATAC-seq reproducible peaks, we selected all the PWM matches assigned with a purity score > 0.7 (the threshold used to define “bound” instances), and then randomly selected 3 times more PWM matches assigned to a purity score < 0.7 (considered as “unbound” instances) to obtain a global set containing 25% / 75% of bound / unbound instances for each TF. The selected regions were extended up to 2kb (1kb in each direction, from the middle of the match), and the 2kb intervals were binned in one hundred 20bp windows. We computed the normalized ChIP-seq and ATAC-seq signal inside each bin. The windows were finally ranked according to the summed ChIP-seq signal in the 10 most central bins (200bp). We finally run a set enrichment analysis with the *fgsea* package to assess whether bound / unbound PWM matches were enriched / depleted along this ranking and computed the enrichment score (ES, positive when bound instances are enriched for highest ChIP-seq signals, negative when unbound instances are depleted for highest ChIP-seq signals) and *p*-values which revealed the strength of the correlation. We performed 1,000 permutations to obtain *p-*values.

### Distribution of transcription factor binding sites in space and time

Using the R package *annotatr* we first created an annotation datasets combining coordinates for hg19 promoters, 3’UTRs, exons, introns and intergenic regions as defined in UCSC, as well as our custom set of enhancers (intersection of the global sets of reproducible H3K4me1 peaks with global sets of reproducible H3K27ac peaks, to focus on enhancers that are active at least in one time point during RAS OIS in WI38 fibroblasts). We then annotated the PIQ binding occurrences for each of the PWM independently, and for the 6 time-points independently. Data were further normalized, to account for differences between time-points in the total number of bound occurrences summed across PWM, and finally they were converted to frequencies. We filtered TFs for which less than 100 binding sites were identified throughout the entire timecourse. TFs were ordered according to the proportions of binding sites located in promoters, introns or exons, and we finally computed the density in migrant, settler and pioneer factors along the ranking.

### Transcription factor co-binding

For every cluster of PWM and time-point independently, we first removed all the bound instances identified outside enhancers. The remaining bound instances for all PWM were then combined for every time point using GEM *regulatory module discovery*^40^ setting at 500 bp the minimal distance for merging nearby TF bound instances into co-binding regions and at 3 the minimum number of TF bound instances in a co-binding region.

#### Global pairwise co-binding heatmap

At this step, we obtained a set of contingency matrices *M_mt_* of dimension *n_mt_* × *j* with *i* the number of co-binding regions for the transcriptomic module *m* at the time point *t* and *j* = 310 PWM clusters, for each time point and each transcriptomic module. We then generated module- and time-specific normalized pairwise co-binding matrices *C_mt_* by computing the normalized cross-product of matrices *M_mt_* defined as:

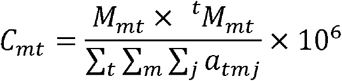

with *a_tmj_* the number of bound instances for the PWM clusters *j*, in transcriptomic module *m*, at the time point *t*. To get a global picture of pairwise co-binding, we summed these matrices and tested for each combination of PWM clusters A and B whether the overlap between bound instances for A and B was significant using a hyper-geometric test defined as:

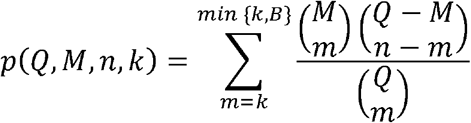

where *Q* is the overall number of regions in the universe, *M* is the number of regions bound by A, *n* is the number of regions bound by B, and *k* the total number of regions bound by A and B. The *p-*values were further corrected for multiple testing using the Bonferroni strategy. We finally clustered the co-binding occurrence matrix using Ward’s aggregation criterion and projected corresponding corrected *q-*values on this clustering.

#### Pair-wise co-binding circos plots

To generate the co-binding circos plots, we used the global time- and, module-specific pair-wise normalized co-binding matrix *C_mt_*described above, after a logarithmic transformation. For each time-point and module independently, we selected the top 500 interactions based on their occurrence *N*. The images were generated using the *Circos* suite^76^.

### Identification of TF regulatory modules

We used the data-sets generated using GEM regulatory module discovery described above. We applied a Hierarchical Dirichlet Process topic model which automatically determines the number of topics from the data, with the hyperparameter for the topic Dirichlet distribution set at 0.1 (encoding the assumption that most of the topics contains a few TFs) and the maximum number of iterations set at 2000. The lexicon usage for each time point and each transcriptomic was explored using a multiple factor analysis (MFA) with the R package *FactoMineR,* and lexicons were further selected based on their goodness of representation on the 3 first components (squared cosine > 0.5).

### TF properties

With the aim of characterizing the binding properties of each TF, we computed the dynamicity, the total number of bound regions, the fraction of bound regions in enhancers and the fraction of bound regions before stimulation.

#### Dynamicity

We quantified the dynamicity of a TF accordingly to the following expression:

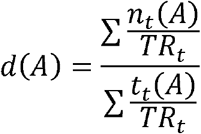

where *d*(*A*) is the dynamicity of TF A; *n_t_*(*A*) is the number of regions bound by A for the first time at time point *t*; *t_t_*(*A*) is the number of regions bound by A at time point *t*, and *TR_t_* is the number of regions bound by any TF in time point *t*,. The factor *TR_t_* was added to the expression to account for differences in the number of reads sequenced by the ATAC-seq protocol and normalizes the number of regions bound by TF A based on the number of bound regions detected at its corresponding time point. Notice that, if all samples have the same amount of TF binding events, this expression is reduced to the quotient of the sum of the regions first bound at each time point by the sum of all regions bound by the TF at each time point. By using this definition, the function *d*(*A*) maps the activity of a TF to the interval 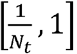, where *N_t_* is the number of time points in the timecourse and is higher as the TF binds to previously not bound regions or leaves already bound regions. In the case of a TF that, for every time point, leaves all its previous bound regions and binds to only regions not previously bound, the numerator will be identical to the denominator, leading to *d*(*A*) = 1. Alternatively, if a TF remains on the same regions it has bound at, *t* = 0, then ∑*n_t_* = *n_0_* and ∑ *n_t_* = *N_t_* * *n_0_*, resulting in 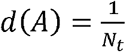. One can observe that, if the same region is bound by TF A in different time points, it will contribute once to the numerator of the expression, while it will contribute to the denominator once for each time point it has been bound to.

#### Total number of bound regions

The number of bound regions was calculated by the following the expression:

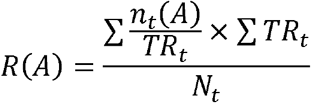

where *R*(*A*) is the normalized number of bound regions by TF A during the timecourse and *n_t_*(*A*), *TR_t_* and *N_t_* are defined as above. The first factor is a normalized sum of the regions bound by TF A, counting each region only once. The second factor scales the result by the mean of the number of regions bound by all TFs on each day.

#### TF percentage of binding at enhancers

The ratio of binding at enhancers, relative to all cis regulatory regions, was assessed by:

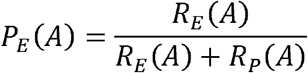

where *P_E_*(*A*) is the percentage of bound regions in enhancers for TF A, *R_E_*(*A*) is the number of regions bound by TF A marked as enhancers and *R_p_*(*A*) is the number of regions bound by TF A marked as promoters.

#### TF prestimulation binding

For each TF, we computed the ratio of regions bound at T_0_, relative to the number of regions bound during the whole timecourse. We used the following definition for the prestimulation binding factor for each TF:

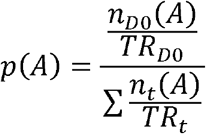

where *p*(*A*) corresponds to the prestimulation binding of TF A and *n_t_*(*A*) and *TR_t_* are defined as above. The numerator of this expression corresponds to the normalized number of regions bound by TF A at t = T_0_, while the denominator is the normalized number of regions bound by TF A during the whole timecourse. Notice the denominator also corresponds to factor *R*(*A*) before scaling.

### Hierarchical transcription factor network

In order to assess the TF chromatin binding hierarchy, i.e. TFs required for the binding of a specific TF, we generated a network for each gene module depicting the precedence of TF chromatin binding. The algorithms mentioned were implemented in R and all networks were visualized in CytoScape^77^.

#### Computing precedence relationships

The edges in the generated networks represent the precedence relationship of TFs: an oriented edge from TF A to TF B, represented as (A, B), means that A was present in at least 30 % of the cis-regulatory regions bound by B at the same instant or before^44^. To account for the difference in the number of reads sequenced for each sample in the ATAC-seq, we normalized the number of regions bound based on the first day they appeared. The weight of an edge from A to B is given by:

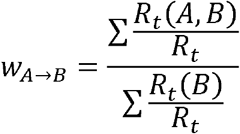

where *R_t_*(*B*) stands for the number of regions first bound by TF B at time point *t*; *R_t_*(*A*, *B*), for the number of regions first bound by TF B at time point *t* that were bound by TF A at time point *i* or before; and *R_t_* represents the total number of regions bound by any TF in time point *t*. In order to handle the networks, we used the *igraph* package^78^.

#### Network simplification

Aiming to analyze the hierarchical relationship of TFs and simplify the interpretation of the network, we performed two operations over each gene module network: Vertex Sort and transitive reduction (TR)^79^. Briefly, the vertex sort algorithm assigns two parameters for each node in the network: the distance, in edges, between the node and the bottom of the network; and the distance between the node and the top of the network. Combined, those parameters allow for the topological ordering of the network, which consists in listing its nodes such that nodes at the top precede downstream nodes. We then defined the ‘top layer’ as the set of nodes with lowest distance to the top of the network, i.e., nodes that have no incoming edges or nodes that assemble a strongly connected component (SCC) with all upstream nodes. Analogously, the ‘bottom layer’ was defined as the set of nodes with lowest distance to the bottom of the network, i.e., nodes with no outgoing edges or that form a SCC with all downstream nodes. The ‘core layer’ comprises nodes that link top layer and bottom layer. Nodes in the core layer that are exactly one edge from both top and bottom layers constitute the ‘single-level core layer’, while nodes that link top and bottom through paths composed of more than one edge form the ‘multi-level core layer’. The result of this procedure for each gene module can be seen in Figure 4 and supplementary data. The TR, in turn, simplifies the network visualization by generating the network with the smallest number of edges that keeps the reachability of the original network.

#### Network validation

We validated our approach by comparing the network produced when applying our method to the ChIP-seq data produced by ^44^ with the network they obtained. Transcription factor ChIP-seq peak files were retrieved from Gene Expression Omnibus (GSE36099, 23 TFs, and 4 time points; note that RUNX1 and ATF4 were discarded from the analysis since one and three time points, respectively, were missing on GEO for those TFs) and preprocessed as previously described to generated time resolved co-binding matrices, further used as an input for our networking algorithm. We computed the precedence relationships among TFs and generated the TF binding hierarchy networks for visualization. We compared the produced TF hierarchy network with the network shown in Figure S4 and in ^44^ using two metrics: sensitivity and specificity. Sensitivity is calculated as the ratio of edges described in this study over the edge number sum for both networks. Specificity is defined as the ratio of the number of edges that were described to not exist in the network produced by our software over the number of edges described to not occur in any of both studies.

#### Proportion of incoming edges based on the classification of the TF source node

Aiming to assess the hierarchy of TFs accordingly to their chromatin dependence and chromatin-opening index, we computed the number of edges connecting the sets of all TFs with a given classification for each gene module. We then divided those values by the number of edges that target TFs with a specific classification. Hence, the proportion of incoming edges based on TF classification is given by:

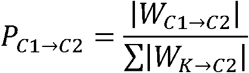

where 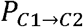 is the proportion of edges from nodes with classification C1 to nodes with classification C2; 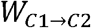 is the set of edges from nodes with classification C1 to nodes with classification C2; *K* can represent either pioneer, settler or migrant and |.| means the cardinality of a set, *i.e.* the number of elements it contains.

We assessed the classification precedence significance for TF interaction with a hyper-geometric test. We consider the sample space as all possible oriented edges in a network with the same number of nodes for each classification as the hierarchy network for a given transcriptional module. Formally:

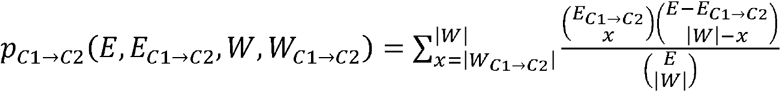

Where *E* is the number of edges on the sample space network, i.e., a fully connected network with the same number of nodes as the TF hierarchy network for a given transcriptional module (excluding self-loops),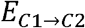 is the number of edges from TFs with classification *C*1to TFs with classification *C*2 in the sample space network, |W|is the number of edges on the TF hierarchy network for a given transcriptional module and 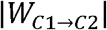 is the number of edges in the same network connecting TFs with classification *C*1 to TFs with classification *C*2.

#### Network visualization

In order to visualize the network, we exported the adjacency matrices in the R environment to CytoScape using the CyREST API^80^. The networks’ layout and style were automated with the help of packages RCy3^81^ and RJSONIO.

### Network mining

With the purpose of identifying key TFs in the transition to the senescent phenotype, we analyzed the TF binding characteristics with their relative location in the chromatin binding hierarchy networks for each gene module. The figures illustrating this analysis were generated with the help of the ggplot2 R package.

#### TF classification

For each network relative to a transcriptional gene module, the number of TF classified as either pioneer, settler or migrant was calculated for each layer, with the subdivision of the core layer as ‘multi-level’ and ‘single-level’ (see “*Network simplification*”). The overrepresentation of TFs with a specific classification in a given layer was evaluated by using a hypergeometric test. We calculated the *p*-value given by:

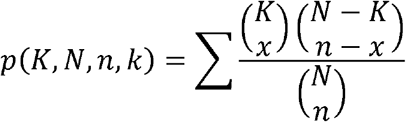

where *K* is the number of TFs with a certain classification in the whole network, *N* is the number of TFs in the network; *n* is the number of TFs that belong to a specific layer and *k* is the number of TFs that belong to the same layer and have the referred classification. The *p*-values were corrected for multiple testing with FDR and a corrected *p* = 0.05 was considered an indicative of enrichment for that specific classification in the corresponding layer.

#### TF dynamicity

For each network relative to a transcriptional gene module, we compared the distribution of the dynamicity of TFs belonging to a certain layer with the distribution of the dynamicity of TFs belonging to the rest of the network. We used the dynamicity index defined previously for each TF, considering only the regions marked as enhancers belonging only to the gene module relative to the network. For each layer in the network, we applied the Kolmogorov-Smirnov test to compare the TF dynamicity distribution for the chosen layer with the dynamicity distribution relative to the TFs belonging to three other layers in the respective network. To account for multiple hypothesis testing, we also performed an FDR correction, considering values of *p* = 0.05 as an indicative of statistical significance.

#### TF number of binding regions

We performed the same analysis as described in the previous section (“*TF dynamicity*”) for the number of bound regions defined in section “*Total number of bound regions*”, instead of the dynamicity index.

#### TF binding characteristics and transcriptional modules

In order to characterize the binding activity of each TF for the different gene modules, we ranked them accordingly to their dynamicity and their number of bound regions. Both parameters for each gene module are shown in Supplementary Figure 4E, which was generated with the *ComplexHeatmap*^82^ and *circlize*^83^ R packages. We used the mean of the ratio dynamicity - number of bound regions to order the TFs. We assessed the significance of pioneer (respectively, migrant) TF enrichment at the top (respectively, bottom) of the ranked clustered list by employing a set enrichment analysis implemented in the package fgsea.

#### TF chromatin binding hierarchy networks overlap

To analyze the similarity between the networks for different transcriptional gene modules, we generated a 7-set Euler diagram, where each set contains the edges present in the TF hierarchy network relative to a gene module. Edges in two different networks are considered equal if they link nodes corresponding to the same TFs in their respective networks. We used the package Vennerable to compute the intersections of all possible network combinations and to create the Euler diagram in Supplementary Figures 4F-I. In this figure, the area of each region is proportional to the number of edges shared by the networks corresponding to the sets that contain the referred region and was calculated using the Chow-Ruskey algorithm^84^. A Euler diagram is similar to a Venn diagram, with the difference that the area of a region representing a set is proportional to the number of elements in the set.

### Analysis of de novo and remnant enhancers

To track combinatorial chromatin state dynamics in space and time, we integrated histone modification ChIP-seq signals at a sub-nucleosomal resolution considering non-overlapping 100bp windows genome-wide using chromstaR (see above), which converts quantitative ChIP-seq data to qualitative chromatin states. For subsequent analysis, since these 100bp windows can be either isolated or organized in stretches experiencing consistent changes in states, we summarized the information at a higher level, and linked them with the histone modification peaks identified using the more classical ChIP-seq and ATAC-seq peak-calling approach (see flowchart). Briefly, after merging all the peaks identified for all the time-points, for all the histone modification and for the ATAC-seq data sets defining *cis-*regulatory regions, we determined the overlap between “poised enhancers“-, “*de novo* enhancers”-, “remnant enhancer” or “constitutive enhancers”-flagged 100bp windows. When an overlap was found, the entire *cis*-regulatory regions were annotated according to the 100bp window it is overlapping with. This operation rendered a list of annotated *cis*-regulatory regions with *de novo*, constitutive, poised or remnant enhancer elements. We finally considered the center +/− 10kb of these elements.

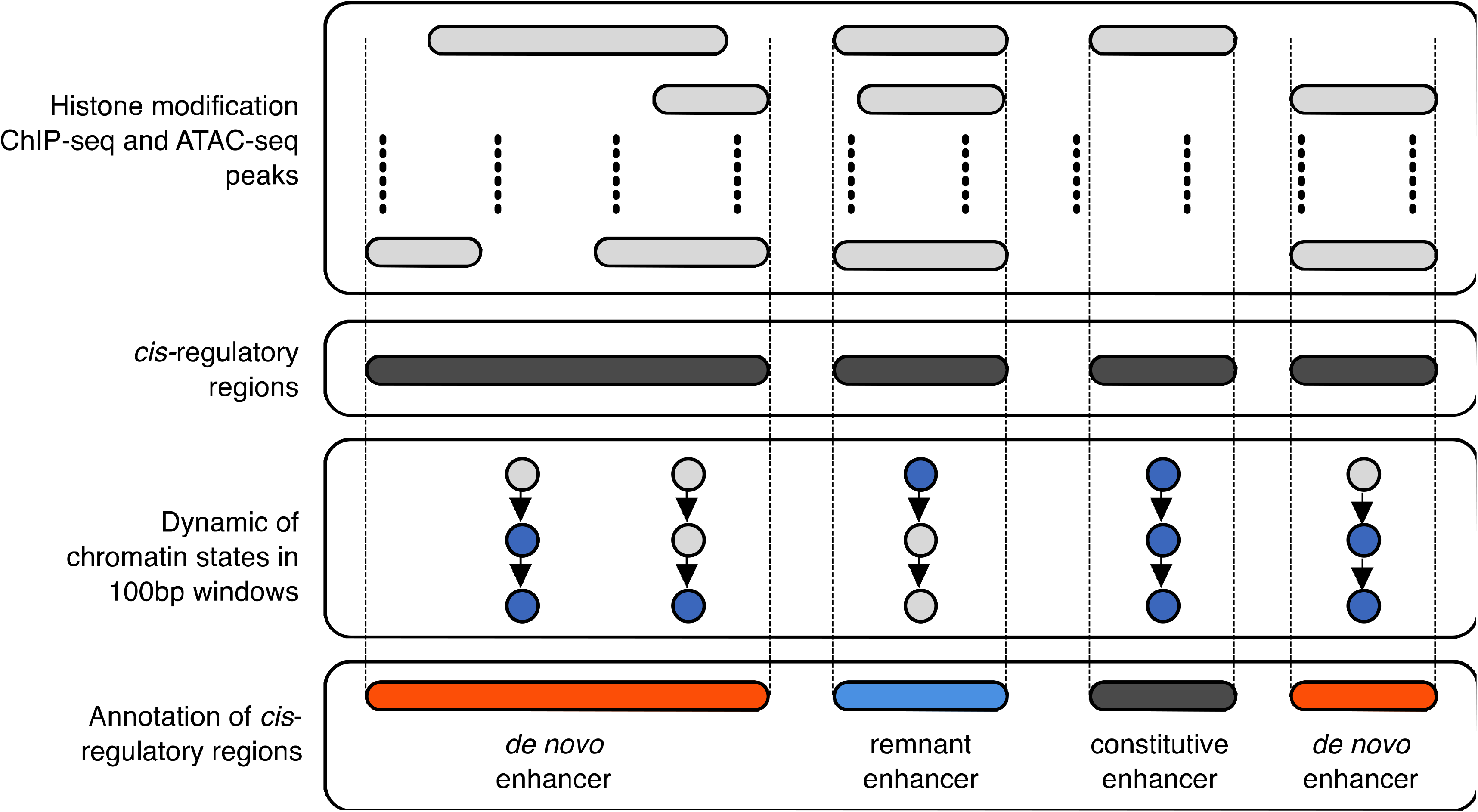

### CRISPR interference (CRISPRi)

hU6-gRNA-hUbc-dCas9-KRAB plasmid was a kind gift from Charles Gerbach (Addgene 71236). gRNA cloning was as published ^34^. Briefly, the plasmid was digested with BsmBI and dephosphorylated before ligation with phosphorylated oligo pairs. The gRNA sequences were listed in the Table 1. The plasmid was then transfected in HEK293T cells, together with packaging plasmids psPAX2 and pMD2.G. 24 hours after fresh medium was added and the medium containing lentivirus was collected and filtered subsequently. Cells were infected for 3 hours. 3 days post infection, cells were passaged and selected with puromycin and used for analyses.

**Table 1.**
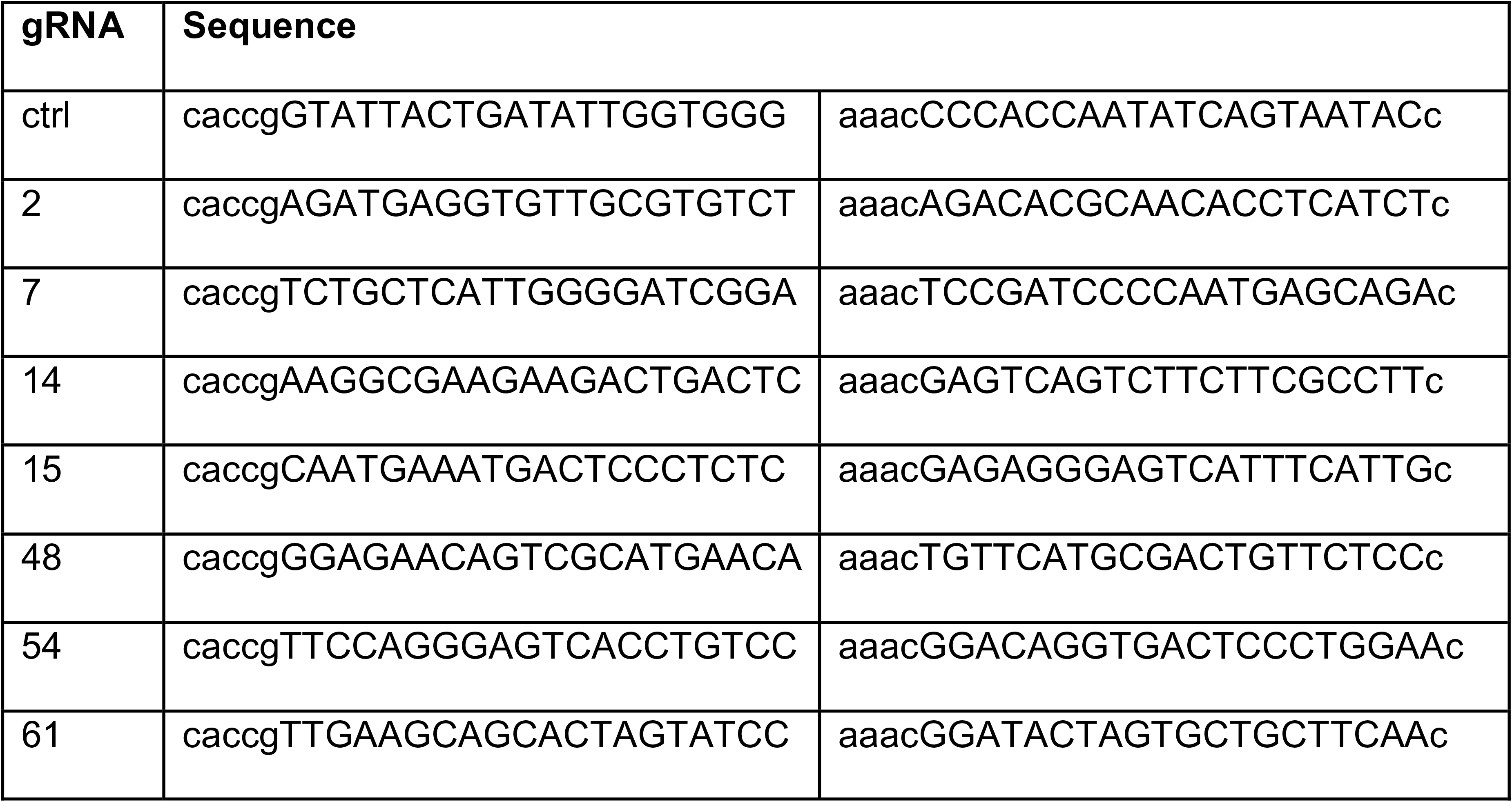
gRNA sequences.

### Immunofluorescence staining and imaging of cells

Immunofluorescence staining was performed as previously published^85^. Cells grown in 96-well plates were fixed with 4% PFA and permeabilised with 0.2% Triton-X in PBS. After blocking, the cells were incubated with primary antibody for 1 hour, and then Alexa Fluor secondary antibody for 30 min. Nuclei were counterstained with DAPI. The antibodies were listed in Table 2. The imaging was carried out by IN Cell Analyzer 2000 (GE Healthcare) with the 20x objective and the quantification was processed using IN Cell Investigator 2.7.3 software.

**Table 2.**
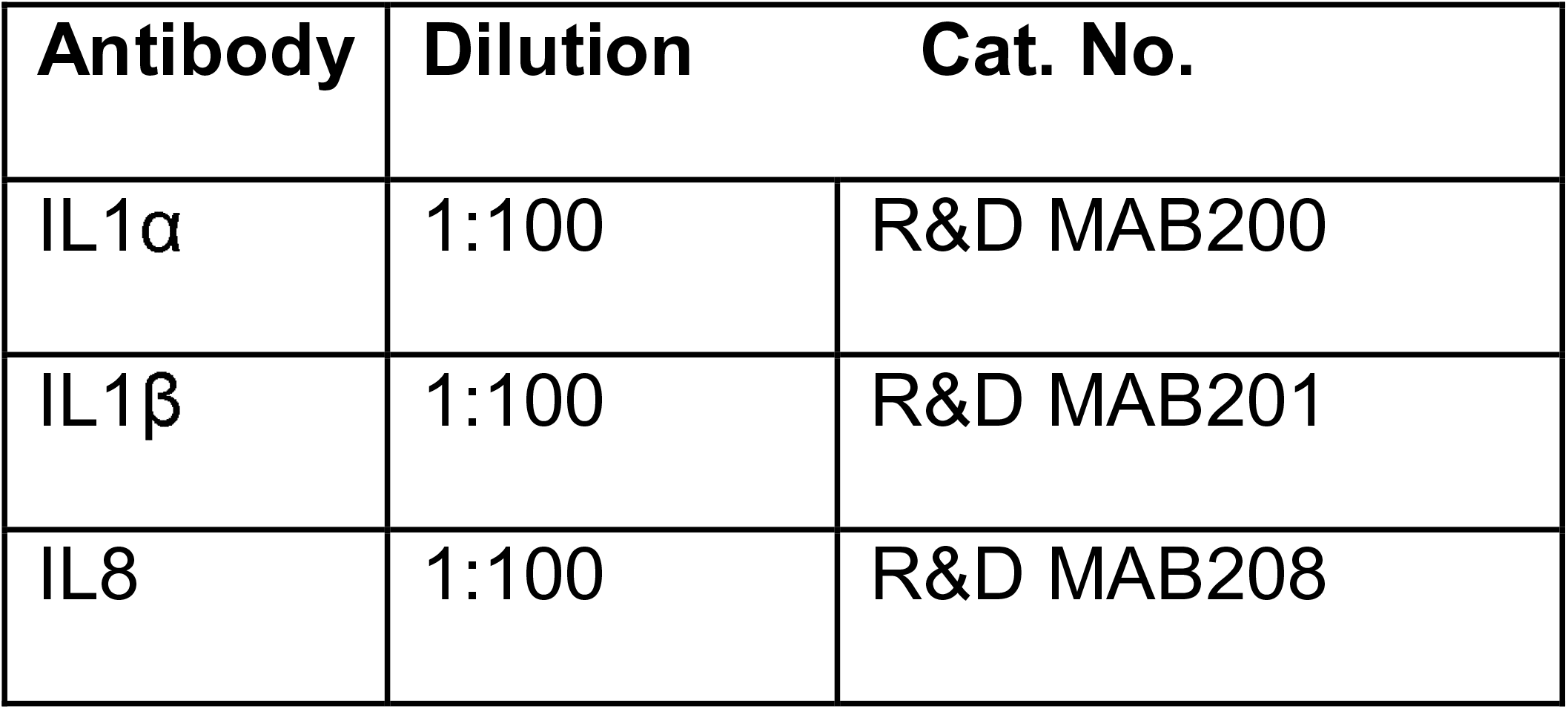
Antibodies.

| Antibody | Dilution | Cat. No. |
| --- | --- | --- |
| IL1α | 1:100 | R&D MAB200 |
| IL1β | 1:100 | R&D MAB201 |
| IL8 | 1:100 | R&D MAB208 |

### Quantitative RT-qPCR

RNA was extracted with TRIzol (Ambion) and RNAeasy Mini Kit (Qiagen) according to the manufacturer’s protocol. Reverse transcription was carried out with SuperScript II RT kit (Invitrogen). Samples were analysed with SYBR Green PCR Master Mix (Applied Biosystems) in CFX96^TM^ Real-Time PCR Detection system (Bio-Rad). Ribosomal protein S14 (RPS14) was used as the housekeeping gene. Primers are listed in Table 3.

**Table 3.**
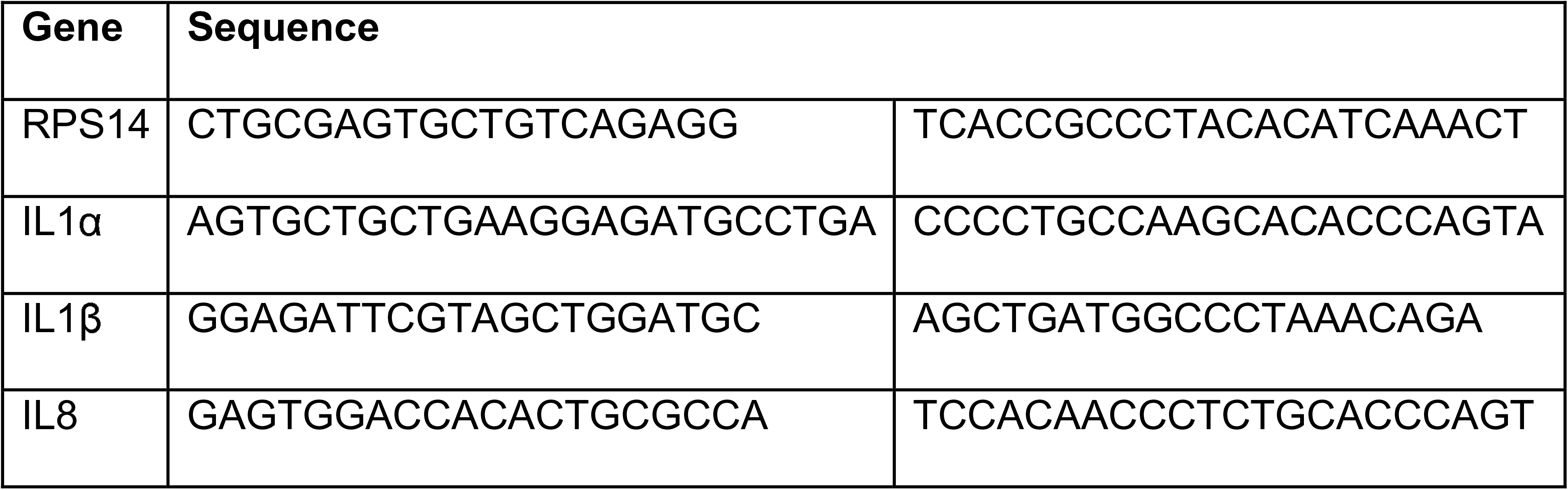
qPCR primer sequence.

### Data availability

All transcriptome data are hosted on Gene Expression Omnibus (BioProject PRJNA439263, accession n°GSE112084). ATAC-seq, and ChIP-seq data (histone modification and transcription factor) are hosted on SRA (BioProject PRJNA439280).

### Code availability for reproducible science

Interactive maps, circus plots, workflows, scripts and software developed to pre-process raw data, perform statistical analyses as well as data mining and integration are available as .html, and R Markdown files provided in Supplementary data hosted on Zenodo (https://zenodo.org, DOI: 10.5281/zenodo.1493872). This archive collapses all the material (including processed data) required to reproduce figures presented in the manuscript.

## ACKNOWLEDGMENTS

We thank all members, in particular Nir Rozenblum, of the O.B. laboratory for fruitful discussions and suggestions through the course of this work. We would like to thank the Transcriptome and Epigenome facility of Institut Pasteur. We thank Claudia Chica for expert advice on ChIP-seq data processing. We thank Ido Amit and Deborah Winter for valuable discussion and technical support. We thank Benno Schwikowski for key insights and technical advice. We also thank Lars Zender, Eric Gilson, and Hinrich Gronemeyer for valuable intellectual input. R.I.M-Z. was supported by La Ligue Nationale Contre le Cancer and was a Mexican National Scientific and Technology Council (CONACYT) and Mexican National Researchers System (SNI) fellow. Lucas Robinson was supported by the Pasteur - Paris University (PPU) International PhD Program and by the Fondation pour la Recherche Médicale (FRM). J.A.N.L.F.F. was supported by La Ligue Nationale Contre le Cancer. O.B was supported by the Pasteur-Weizmann Foundation, ANR-BMFT, Fondation ARC pour la recherche sur le Cancer, La Ligue Nationale Contre le Cancer, INSERM-AGEMED. Research reported in this publication was supported by the National Cancer Institute of the National Institutes of Health under Award Number R01CA136533. The content is solely the responsibility of the authors and does not necessarily represent the official views of the National Institutes of Health. O.B. is a CNRS Research Director DR2.

## AUTHOR CONTRIBUTIONS

R.I.M-Z, P.-F.R and O.B conceived of the study and conceptual ideas. R.I.M-Z, P.-F.R and O.B. planned, designed experiments, interpreted data and wrote the manuscript. All authors discussed the results and contributed to the final manuscript. R.I.M-Z generated the cell culture system and performed ChIP-seq, ATAC-seq and RNAi experiments. P-F.R performed computational analyses, designed bioinformatics pipelines and prepared figures. L.R. performed senescence characterization studies and performed ChIP-seq experiments. J.A.N.L.F.F. generated TF networks. G.D. generated Affymetrix microarray data. B.S. and J.G. performed CRISPRi experiments. U.H. supported the study. O.B. supervised, managed and obtained funding for the study.

## DECLARATION OF INTERESTS

The authors declare no conflict of interest.

---

**Figure S1:**
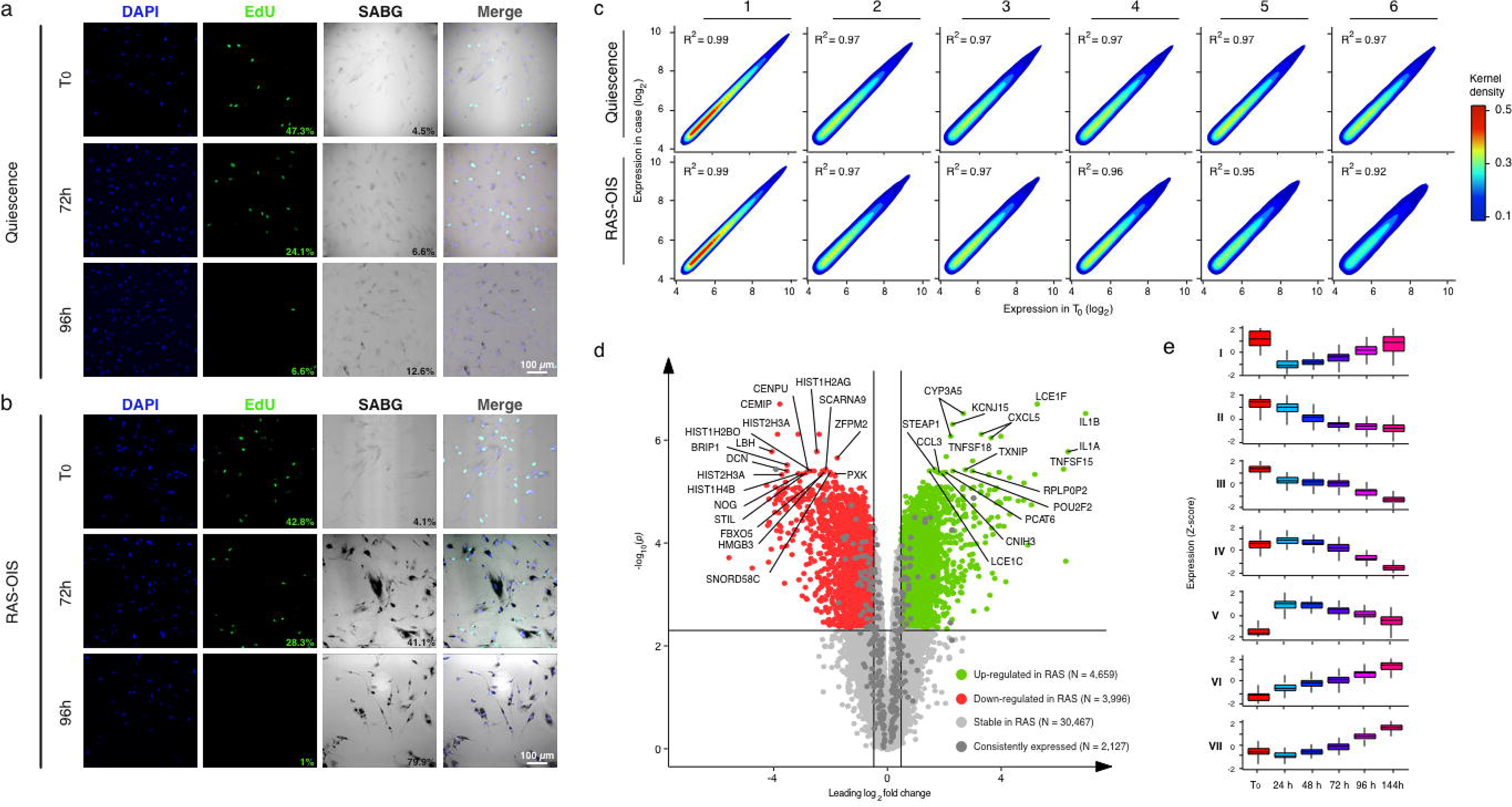
Multi-state establishment of the senescence transcriptional program. **(a-b)** Characterization of quiescence and RAS-OIS cells. **(a)** Representative DAPI, EdU, SABG (from left to right) indirect fluorescence and phase contrast microscopy images of WI38 fibroblasts undergoing quiescence at indicated time-points. Proliferative capacity is % of EdU-positive staining cells. Scale bar, 100µm. **(b)** Representative DAPI, EdU, SABG (from left to right) indirect fluorescence and phase contrast microscopy images of WI38 fibroblasts undergoing OIS at indicated time-points. Scale bar, 100µm. **(c)** Distribution of gene expression levels as kernel density estimates for time-resolved quiescence and RAS-OIS transcriptomes. Pearson’s correlation coefficient (R^2^) is shown. **(d)** Volcano plot of RAS-OIS time-series transcriptome data. Dark grey dots highlight genes sharing a common gene expression pattern between quiescence and OIS time-series experiments and were removed to define the RAS-OIS specific temporal transcriptomic signature used for all further downstream analyses. **(e)** Boxplot depicting expression patter for each of the RAS-OIS transcriptomic modules. Data are expressed as row Z-score.

**Figure S2:**
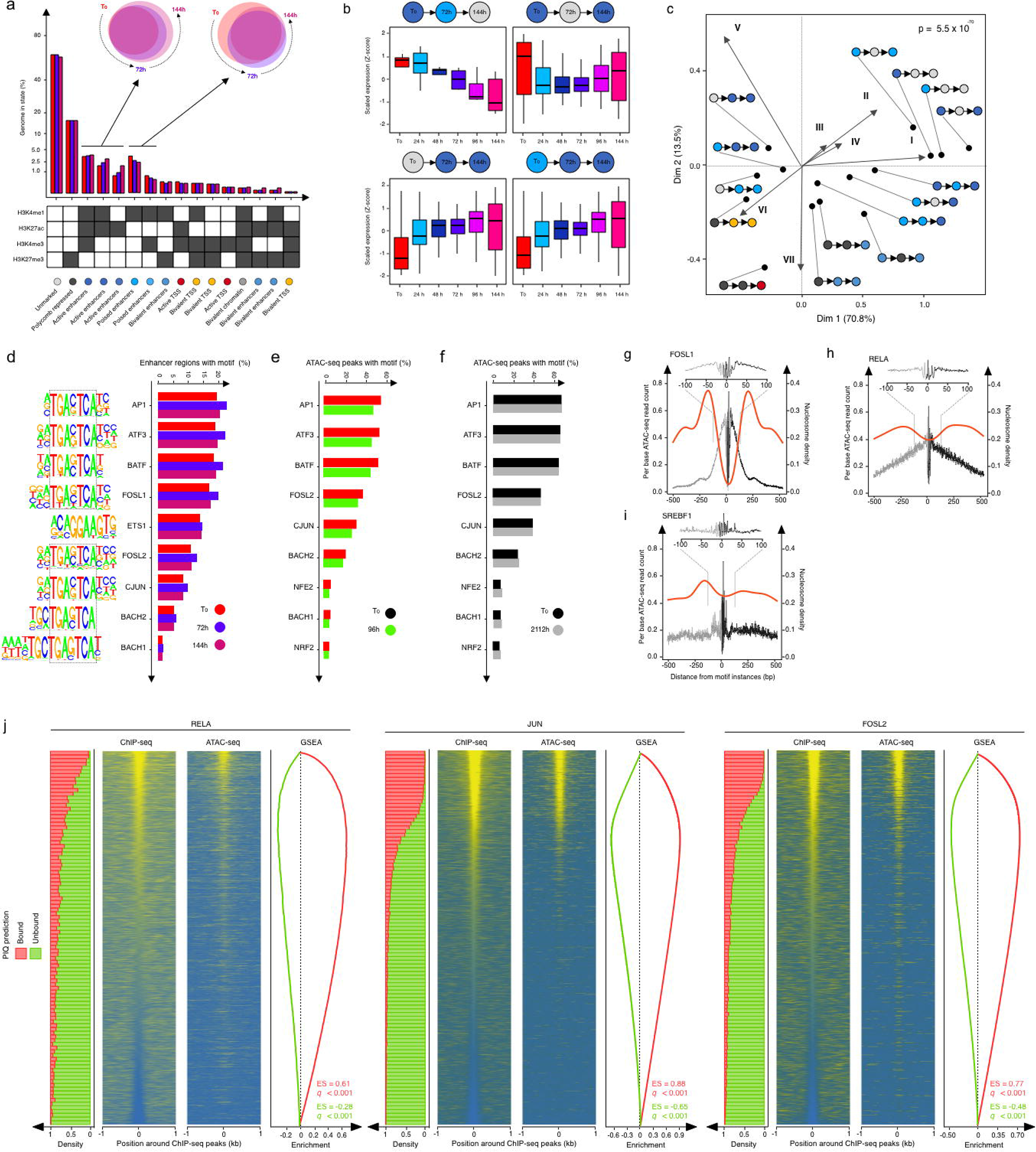
A dynamic enhancer program shapes the senescence transcriptome. **(a)** Histogram showing the percentage of genome covered by each chromatin state at indicated time. Bottom table assigns histone modification combinations (grey: presence, white: absence) to biologically meaningful mnemonics. Venn diagrams highlight the specificities and overlaps in chromatin states associated with active (left) and poised enhancers (right) at indicated time-points. **(b)** Boxplots showing the distribution of relative gene expression (row Z-score) through time for genes associated to regions undergoing different chromatin state changes. The pictogram at the top of each graph describes the class of chromatin state change considered. **(c)** Asymmetric biplot for correspondence analysis between changes in chromatin states and gene expression modules. The *p-*value reflects the strength of the association using a χ^2^ test. Only the top 20 contributing and best projected (squared cosine > 0.5) chromatin state changes are shown. **(d-f)** Most enriched sequence motifs in **(d)** active enhancers or **(e-f)** ATAC-seq peaks at each time point for the **(d)** RAS-OIS, **(e)** replicative senescence, and **(f)** quiescence time courses. **(d)** Motif logos are shown on left of the histogram. Black, dotted boxes highlight the core motif for AP1 transcription factor family members. Note that the transcriptional repressor BACH shares this motif. **(g-i)** ATAC-seq (grey lines for forward, black lines for reverse reads) and nucleosome (red line) footprints for **(g)** AP-1 FOSL1 (pioneer), **(h)** RELA (settler), and **(i)** SREBF1 (migrant). **(j)** Comparison between PIQ predictions and RELA (left), AP-1-JUN (middle) and AP-1-FOSL2 (right) ChIP-seq. The two density heatmaps at the center of each panel illustrate ChIP-seq (left) and ATAC-seq (right) signals computed in 10bp non-overlapping windows at selected bound- (25%) and unbound- (75%) predicted PWM hits ± 1kb ranked according to the ChIP-seq signal in the most central 100bp. The stack histogram on the left shows the distribution of bound (red) and unbound (green) PWM hits as defined by PIQ along the ranking. The curves on the right depict the evolution of the enrichment score (ES) along the ranking as defined with a set enrichment analyses (SEA) comparing the ChIP-seq signal and the bound (red) and unbound (green) status of the PWM hit. For each SEA, we performed 1 000 permutations and provide the associated Benjamini–Hochberg adjusted *p*-value and ES score.

**Figure S3:**
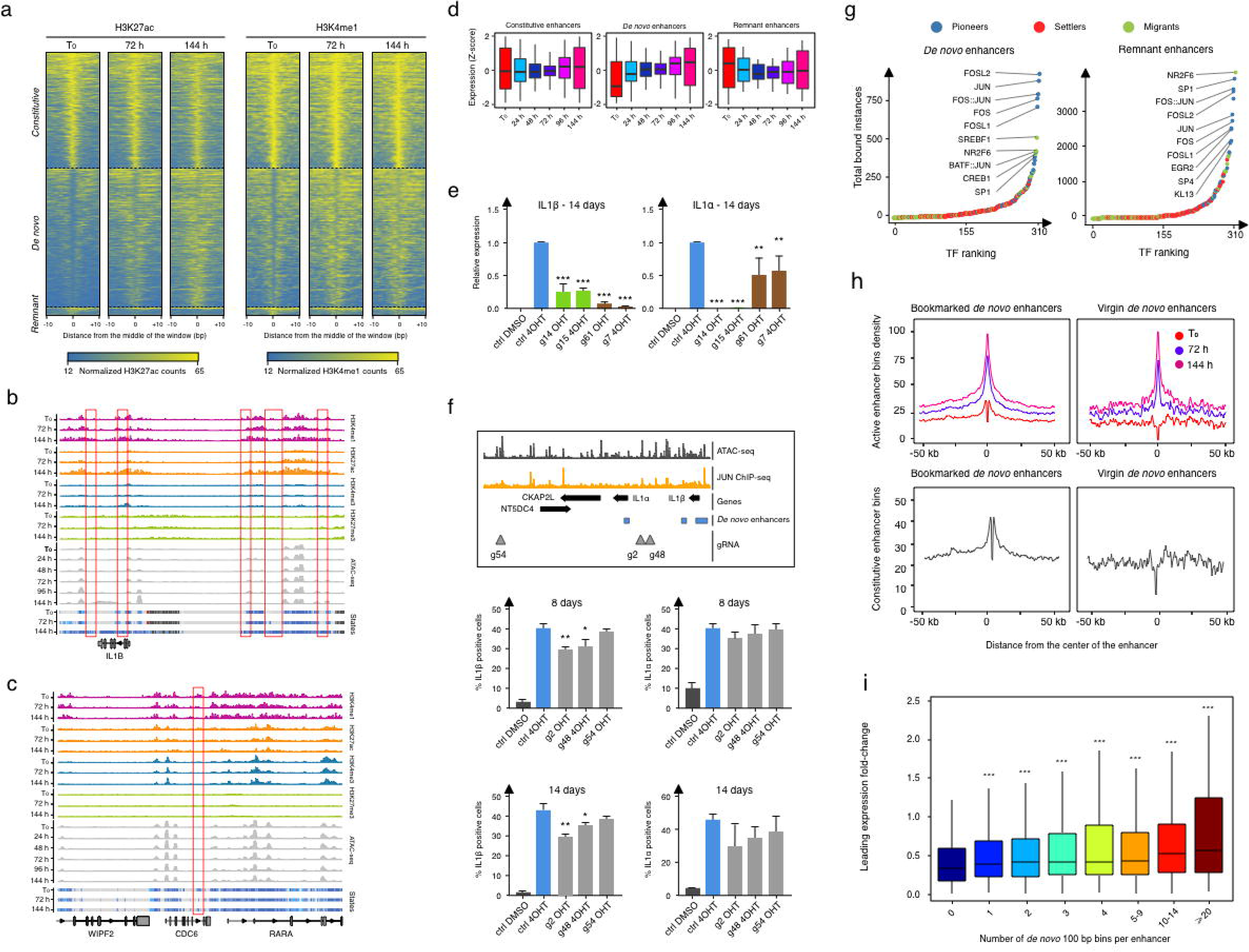
AP-1 pioneer TF bookmarking of senescence enhancer landscape foreshadows the senescence transcriptional program. **(a)** Density heatmaps of normalized H3K27ac and H3K4me1 ChIP-seq signals computed in 10bp non-overlapping windows at enhancers +/− 10kb grouped by enhancer status (constitutive, *de novo* or remnant) at indicated time-points after RAS-OIS induction. **(b-c)** Representative genome browser screenshots of normalized H3K4me1 (pink), H3K27ac (orange), H3K4me3 (blue) and H3K27me3 (green) ChIP-seq and ATAC-seq (light grey) profiles and chromatin states at **(b)** *IL1ß* and **(c)** *CDC6* gene loci. Red boxes single-out **(b)** IL1ß *de novo* and **(c)** CDC6 remnant enhancers. **(d)** Boxplots depicting the distribution of relative gene expression (row Z-score) through time for genes associated with constitutive (left), *de novo* (middle) and remnant (left) enhancer windows. **(e)** RAS-OIS cells at day 14 infected with dCas9-KRAB and individual guides (g14, g15, g61, and g7) and analyzed by RT-qPCR for the expression of IL1α or IL1β as described in Figure 3b. Data represent mean ± SD (n=3). *p<0.05, ***p<0.001. Comparison with ctrl 4OHT, one-way ANOVA (Dunnett’s test). **(f)** RAS-OIS cells were infected with dCas9-KRAB and individual guides (g2, g48 and g54) for non-enhancer regions (outside *de novo* enhancers) as described in Figure 3b. 8 or 14 days after infection, cells were stained for IL1α or IL1β by indirect immunofluorescence and percentage positive cells were quantified (n=3 for 8 days and n=2 for 14 days). Data represent mean ± SD. *p<0.05, **p<0.01, ***p<0.001. Comparison with ctrl 4OHT, one-way ANOVA (Dunnett’s test). **(g)** Rank plot depicting the summed occurrences for TFs binding in proliferating cells (T_0_) in *de novo* enhancers (left) and after replicative senescence in remnant enhancers (right). Top ten TFs are highlighted. **(h)** Metaprofiles showing the density in “active enhancer”-flagged genomic bins (top) and “constitutive enhancer”-flagged genomic bins (bottom) in the vicinity (+/− 50kb) of TF bookmarked *de novo* (left) and TF virgin *de novo* enhancers (right). The density in “active enhancer”-flagged genomic bins is provided for the indicated time points. **(i)** Boxplot showing the correlation between absolute leading log_2_ expression fold-change and the number of genomic bins flagged as “*de novo”* enhancers per enhancer. ***: *p*-value < 10^−3^, Student’s *t*-test considering regions with 0 “*de novo”* enhancers bins as a control.

**Figure S4:**
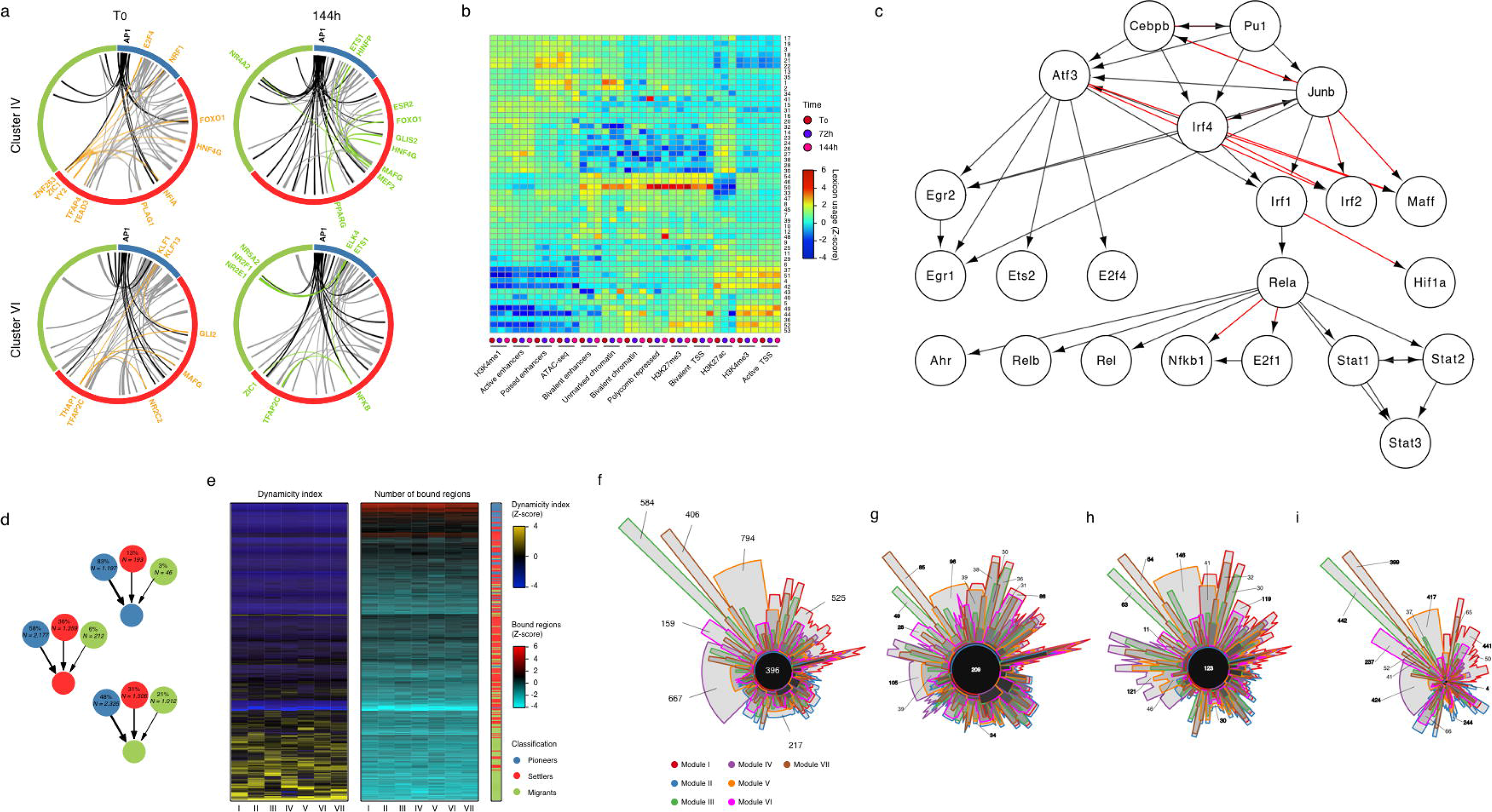
A hierarchical TF network defines the senescence transcriptional program. **(a)** Representative circos plots summarizing pairwise transcription factor co-binding at enhancers for down-regulated transcriptomic module (IV, top) and up-regulated transcriptomic module (VI, bottom) at indicated time-points. Co-interactions involving AP1 are shown in black. Selected examples of gained (green) and lost (orange) interactions are highlighted. Pioneer TFs blue, settler TFs red, migrant TFs green. See also dynamic circos plot movies in Supplementary Data (see under Code availability in Material and Methods). **(b)** Heatmap showing the overlap between TF lexicons (rows) and chromatin states, ChIP-seq and ATAC-seq peaks (columns). The dendrograms were computed by applying hierarchical clustering on the fraction matrix with Pearson’s correlation and average linkage. **(c)** Validation of TF network algorithm using TF ChIP-seq published data sets (Garber et al., 2012). Edges colored in gray were detected in both studies and edges colored in red were found only the analysis performed by Garber et al.. The displayed edge set is the same as in Figure 5A (Garber et al., 2012). We employed a transitive reduction step in order to facilitate visualization. Comparison of the two networks resulted in a sensitivity of 88,9 % and a specificity of 100 %. **(d)** Ratio of incoming edges based on the classification of the TF source node. The relative and absolute number of edges corresponding to all seven modules are displayed inside the nodes, which are colored accordingly to TF classification as in previous panels. The thickness of links is proportional to the relative number of TF hierarchy edges connecting nodes with the corresponding classification. **(e)** Number of bound regions and dynamicity index for each TF (rows) across all gene modules (columns). The left heatmap depicts the dynamicity index scaled by column. The middle heatmap depicts the square root of number of bound regions scaled by column. The right single-column heatmap illustrates TF classification. **(f)** Venn diagram showing specificities and overlaps of TF interactions in each gene module. Each set corresponds to the TF-TF network edges identified for a given transcriptomic module. The global area of each set is proportional to the number of edges in its respective transcriptomic module and was calculated with the Chow-Ruskey algorithm. **(g-i)** Chow-Ruskey diagrams for edges **(g)** originating only from TFs at the top of hierarchy, **(h)** connecting only TFs at the core layer or **(i)** reaching only TFs at the bottom. Note that edges at the top of the hierarchy are shared among the gene modules while edges towards the bottom of the hierarchy are module-specific.

**Figure S5:**
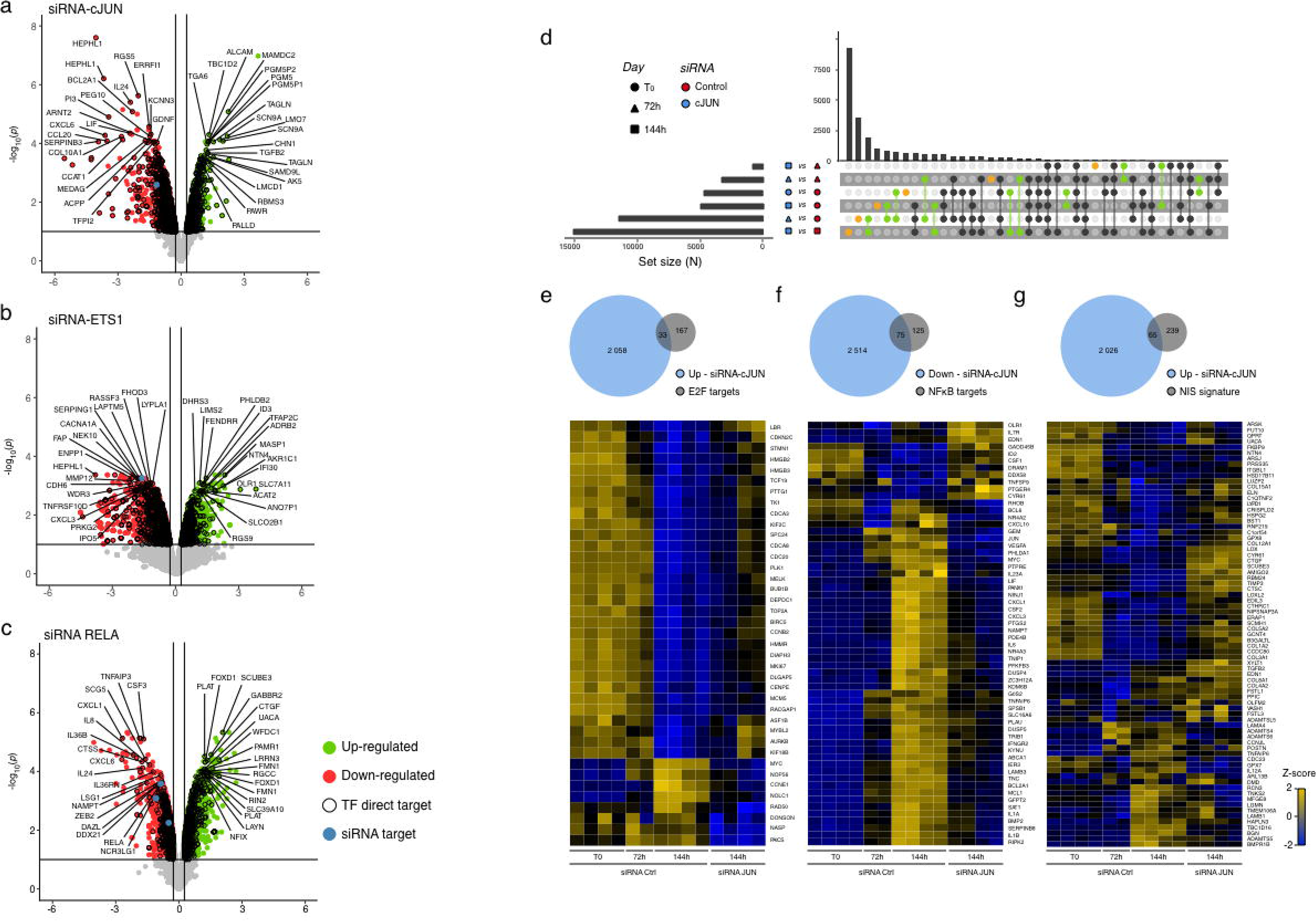
Hierarchy Matters: Functional Perturbation of AP-1 pioneer TF, but no other TF, reverts the senescence clock. **(a-c)** Volcano plots depicting the −log_10_ *p*-value as a function of the log_2_ fold-change in gene expression defined by a differential analysis conducted with *limma* to highlight the effect of siRNA-mediated **(a)** AP-1-c*JUN*, **(b)** *ETS*1 and **(c)** *RELA* depletion in senescent RAS-OIS cells at day 6 (144h). Blue dots in respective plots indicate probes corresponding to AP-1-c*JUN*, *ETS*1 and *RELA*. Black outlined dots highlight direct targets of AP-1-c*JUN*, *ETS*1 and *RELA*. **(d)** Upset plot depicting specificities and overlaps in differentially expressed genes of siRNA-Control and siRNA-*JUN* silenced OIS fibroblasts at indicated time-points. The yellow dots highlight gene sets specific to a single comparison set, while green dots highlight gene sets find in two different pair-wise comparison. **(e-g)** Venn diagrams (top) and heatmaps (bottom) depicting the overlap between genes belonging to **(e)** E2F-, **(f)** NFκB target, and **(g)** N1ICD-induced senescence (NIS) gene signatures. Venn diagrams show the overlap of up-regulated genes after siRNA-mediated AP-1-c*JUN* knock-down for upregulated E2F- (*i.e.* pro-proliferation genes), NIS- (*i.e.* early SASP genes), and downregulated NFκB target genes (*i.e.* late SASP genes) RAS-OIS cells at day 6 (144h). Bottom heatmaps show the comparison of gene expression profiles of siRNA-Control (siCtrl) and siRNA-cJUN treated cells undergoing RAS-OIS at indicated time-points. Data are expressed as row Z-score. E2F targets and NFκB targets were defined according to Molecular Signature Database (MSigDB).

